# Conformational plasticity of mitochondrial VDAC2 controls the kinetics of its interaction with cytosolic proteins

**DOI:** 10.1101/2024.11.08.622687

**Authors:** William M. Rosencrans, Harisankar Khuntia, Motahareh Ghahari Larimi, Radhakrishnan Mahalakshmi, Tsyr-Yan Dharma Yu, Sergey M. Bezrukov, Tatiana K. Rostovtseva

## Abstract

The Voltage Dependent Anion Channel (VDAC) is the major conduit of water-soluble metabolites and small ions into and out of the mitochondria. In mammals, VDAC exists in three isoforms, VDAC1, VDAC2, and VDAC3, each characterized by distinct tissue-dependent distribution and physiological role. VDAC2 is the most notable among the three isoforms because its knockout results in embryonic lethality and regulates the BAK/BAX-dependent apoptosis pathways. Yet, understanding of the biophysical underpinnings of VDAC2 functions remains limited. In this study, we reevaluate VDAC2’s properties, utilizing recombinant human VDAC2 WT and its three mutants – cysteine-less VDAC2, VDAC2 with truncated first 11 residues, and E84A - to explore the biophysical basis that distinguishes VDAC2 from the other isoforms using single-molecule electrophysiology. We found that contrary to VDAC1 and VDAC3, which are characterized by a unique open state, VDAC2 displays dynamic switching between a few high-conductive anion-selective substates. We employed α-synuclein (αSyn) – a known potent cytosolic regulator of VDAC1 and VDAC3 – as a sensitive molecular probe to show that it induces characteristic blockage events in all open substates of VDAC2 but with up to ten-fold different on-rates and blockage times. A substate with higher conductance always corresponds to a higher on-rate of the αSyn-VDAC2 interaction but proportionally lower blockage times. This gives the same equilibrium constant for all substates, thus resulting in the same affinity of the αSyn-VDAC2 interaction. The pronounced difference is limited to the kinetic parameters, suggesting that once the αSyn molecule is captured, its physical state and free energy are the same for all substates. These striking results imply that the αSyn molecule senses the dynamic structural variations within the channel prior to its final capture by the pore. We propose that the discovered conformational flexibility may allow VDAC2 to recognize a larger number of binding partners, thus explaining the physiological significance of this isoform, namely, its ability to adapt to mitochondrial metabolic conditions in cells dynamically.

## Introduction

The Voltage Dependent Anion Channel (VDAC) is the most abundant protein in the mitochondrial outer membrane (MOM) which represents a class of β-barrel channels originally derived from the endosymbiotic bacterial ancestors of modern mitochondria. It is the major pathway for water-soluble metabolites and small ions to cross the MOM. In mammals, there are three VDAC isoforms: VDAC1, VDAC2, and VDAC3. Despite ∼70% sequence similarity between the isoforms and the ability of them all to form large conductive anionic channels (∼4 nS in 1M KCl at room temperature), which gate in response to the applied voltage when reconstituted into a planar lipid membrane (PLM), each VDAC isoform has a distinct physiologic role. VDAC1 and VDAC2 are the most abundant isoforms, with VDAC3 expressed in low levels (∼ 10 % of the total VDACs) in most tissues, except in the testis [1,2]. VDAC2 is the most expressed isoform in brain tissue, as well as in placental endothelium [3]. Studies in mice demonstrate that VDAC1 or VDAC3 knockouts are not lethal but result in metabolic impairment in the case of VDAC1 [4] and male infertility in the case of VDAC3 knockouts [5]. VDAC2 knockout results in embryonic lethality or severely diseased neonatal pups [6,7]. VDAC2 alone has been shown to regulate the BAK/BAX-dependent apoptosis pathways, rationalizing the results of the murine studies [6–8]. In this paradigm, apart from its channel activity, VDAC2 acts as a membrane anchor for BAK/BAX within the MOM. In the case of BAK, binding to VDAC2 impairs its oligomerization in the MOM and subsequent apoptosis via cytochrome c release [6,9]. Studies on the biophysical basis of VDAC2’s unique physiology have been complicated by the fact that, while multiple high-resolution structures of VDAC1 exist, human VDAC2 has eluded structural determination. VDAC2 differs from VDAC1 most obviously in an 11-residue N-terminal extension (NTE) and the presence of nine cysteines compared to two in VDAC1 [10]. Naghdi et al. mutagenesis studies [8] showed that neither the NTE nor the extra cysteines were essential for the VDAC2-BAK interaction, instead finding that a cytosol-facing loop connecting two beta strands was the site of the interaction. Biophysical studies on VDAC2 have demonstrated that the NTE was important for maintaining the stability of the channel, compensating for the increased number of cysteines [11]. VDAC2 channel properties are superficially identical to VDAC1 [11–14], however studies on VDAC2 electrophysiology have identified subtle, yet unexplored, differences. The first study systemically comparing the channel properties of each VDAC isoform [12], found that VDAC2 conductance deviated from a unimodal gaussian distribution characteristic for VDAC1 or VDAC3. The authors speculated that VDAC2 existed in at least two states with different anionic selectivities, the finding recapitulated in more recent studies [14]. VDAC2 has been shown to be slightly more permeable to Ca^2+^ than VDAC1 [15]. The structural features and physiologic role of these states remain unidentified.

VDACs are known to be regulated by cytosolic proteins, including α-Synuclein (αSyn), an intrinsically disordered neuronal protein intimately associated with Parkinson’s disease (PD) [16,17]. The large aggregates and fibrils of αSyn found in Lewy bodies in the brains of PD patients, are a hallmark of PD [18]. αSyn aggregation and oligomerization are associated with its increased expression in dopaminergic neurons in PD and are generally considered to be pathological phenomena [18,19]. However, the exact mechanism of αSyn-induced pathology remains mainly unknown. Especially the role of monomeric αSyn continues to be poorly understood. αSyn was found to be associated with both mitochondrial membranes, causing impairment of the mitochondrial respiratory complexes [20,21] [22], oxidative stress [23], and fission [24] [25]. At the MOM, monomeric αSyn interacts with VDAC (reviewed in [26]) and translocates through VDAC1 into the mitochondrial intermembrane space as was shown in neuroblastoma and HeLa cells [27,28].

In vitro, monomeric αSyn at nanomolar concentrations induces voltage-dependent reversible blockages of reconstituted VDAC channel conductance, seen as rapid current fluctuations on the scale of 1-100 ms between the VDAC open and αSyn-blocked state with the residual conductance of ∼0.4 of the open state [29]. αSyn is a 140-residue polypeptide which consists of three well-defined domains: an N-terminus amphipathic membrane binding domain, a central non-polar NAC domain, and a disordered highly negatively charged C-terminal domain [30]. αSyn is essentially disordered in solutions, but upon binding to the anionic lipid membranes, its N-terminal domain adopts different α-helical conformations while leaving the C-terminus disordered above the membrane surface [30–32]. Based on these structural data and the results of extensive kinetic analysis of VDAC interaction with αSyn at the single-channel level, the current model is that the N-terminal domain of αSyn remains membrane-bound, and the anionic C-terminal domain is driven by the applied negative potential into the VDAC pore [33]. At lower applied voltages, it retracts back to the same membrane surface after residing in the pore for some time which depends on the voltage magnitude [33]. The increasing negative voltage keeps the anionic C-terminus inside the pore longer, which is seen as an exponential voltage dependence of the blockage time. The blocked state displays the reversal of selectivity, going to cationic versus anionic selectivity of the open state [34]. At higher applied transmembrane voltages (|*V*| > 35-45 mV depending on experimental conditions), the blockage time starts to decrease with voltage, which is explained by αSyn translocation through the VDAC pore to the opposite side of the membrane [29]. The translocation regime of αSyn was confirmed by direct experiments enabling real-time monitoring of a single αSyn molecule translocating through the VDAC pore [34].

In light of the proposed model, we can suggest a few immediate physiological implications of VDAC-αSyn interaction, driving regulation of ATP/ADP and Ca^2+^ fluxes across MOM. When the highly negatively charged C-terminal tail of αSyn is captured by the VDAC pore, it creates an electrostatic barrier for ATP/ADP [35] and, by inversing pore selectivity, promotes Ca^2+^ transport [15,36]. The increased mitochondrial Ca^2+^ uptake in HeLa cells in response to αSyn overexpression [37] provides in vivo support to these predictions. Another anticipated implication of the proposed mechanism is the ability of αSyn to translocate through VDAC. Translocation of αSyn into mitochondrial intermembrane space was recently confirmed in neuroblastoma and HeLa cells [27,28]. By entering mitochondria, αSyn could target electron transport chain (ETC) complexes in the inner membrane, thus inducing mitochondrial dysfunction and eventually leading to cell death [22].

The structures of VDAC2 and VDAC3 are not available yet, but in silico reconstructions, based on a high sequence similarity between all three isoforms, predict their structures to be almost identical to VDAC1, which consists of 19 β-strands and an α-helical N-terminus lying in the middle of the pore [38]. The position of NTE in VDAC2 remains unknown. Considering similar β-barrels formed by all three VDAC isoforms and the proposed model of αSyn-VDAC interaction, one would expect a similar interaction of αSyn with all isoforms. However, it was found that the rate of αSyn capture by VDAC1 (the on-rate) is 100 times higher than that by VDAC3 [39] thus demonstrating, on a single-channel level, a clear quantitative difference between these two isoforms. Furthermore, the voltage required for αSyn to translocate through VDAC3 appeared to be 12 mV smaller than that for VDAC1 [39]. These data enabled us to speculate that the difference in interaction with known, such as αSyn and dimeric tubulin, and still unidentified VDAC regulators could constitute a general mechanism that distinguishes isoforms in vivo [40]. Therefore, we reasoned that VDAC2 may differ from the other two isoforms by its interaction with cytosolic regulator proteins thus providing a biophysical basis for its unique physiological role.

In this study, we reevaluate VDAC2’s properties at the single-channel level. We utilize recombinant human VDAC2 WT and its three mutants to understand the biophysical basis that distinguishes VDAC2 from the other two isoforms. We found that contrary to VDAC1 and VDAC3, which are characterized by a unique high conductive or “open” state, VDAC2 is a dynamic channel that spontaneously switches between multiple high-conductive open substates that vary by conductance and occurrence while remaining anion selective – a defining feature of the ATP-permeable open states of VDAC. Importantly, we show that monomeric αSyn induces similar characteristic blockage events in all VDAC2 open substates but with up to ten-fold different on-rates and blockage times. The higher on-rate of the αSyn-VDAC2 interaction always corresponds to the substate with the higher conductance when measured for the same single channel. However, the equilibrium constant of the αSyn-VDAC interaction, which takes into account both the on-rates and the blockage times, remains the same for all substates, independently of their conductance. The finding that the pronounced difference is limited to the kinetic parameters only suggests that once the αSyn molecule is captured, its physical state and free energy in the pore are the same for all substates.

We propose that the appearance of distinct substates within the same channel and their different interaction kinetics with αSyn reveal the dynamic plasticity of VDAC2, which suggests a key for explanation of the exceptional role of this multifaceted channel in the cell. Indeed, the discovered conformational flexibility may allow VDAC2 to recognize a larger number of binding partners. This data could tentatively explain the physiological significance of VDAC2: its ability to dynamically adapt to metabolic cell conditions and change the rates of interaction with its multiple cytosolic protein partners.

## Materials and Methods

### Cloning, recombinant protein production, and purification

Recombinant VDAC2 WT and VDAC2 E84A were purified and isolated as described in *Supplemental Methods* and stored in the buffer containing 25 mM HEPES pH 7.4, 100 mM NaCl, 0.1% LDAO, 1mM TCEP, and 1mM EDTA. The cysteine-less VDAC2 mutant (VDAC2-ΔCys) and VDAC2 mutant without N-terminal extension residues 1-11 (VDAC2-ΔNE) were purified and isolated as previously describes [11]. Recombinant zebrafish VDAC2 (zfVDAC2) was the generous gift of Dr. Johann Schredelseker (Walther Straub Institute of Pharmacology and Toxicology, Munich, Germany). Recombinant αSyn WT was the generous gift of Dr. Jennifer Lee (NHLBI, NIH). αSyn was purified and characterized as described previously [41] and stored at −80^0^C.

### VDAC reconstitution and conductance measurements

The procedure of VDAC reconstitution into lipid bilayers was described previously [29,42] and in *Supplemental methods*. Planar lipid membranes (PLMs) were formed from lipid mixtures of dioleoyl-phosphatidylglycerol (DOPG), dioleoyl-phosphatidylcholine (DOPC), and dioleoyl-phosphatidylethanolamine (DOPE) in a ratio of 2:1:1 (wt/wt) in single-channel experiments or from soybean polar lipid extract (PLE) in multichannel gating experiments. All lipids were purchased from Avanti Polar Lipids. Typically, channel insertion was achieved 2–20 min after the addition of 0.2–0.5 μl of VDAC2 diluted in 2.5% Triton X-100 buffer (50 mM KCl, 100 mM Tris at pH 3.35, 1 mM EDTA, 15% (vol/vol) DMSO, 2.5 % Triton X-100) on the Teflon partition facing the *cis* compartment. The potential is defined as positive when it is greater at the side of VDAC addition (*cis* side). Current recordings were performed as described previously [29] using an Axopatch 200B amplifier (Axon Instruments) in the voltage-clamp mode. Data were filtered by a low-pass 8-pole Butterworth filter (Model 900 Frequency Active Filter, Frequency Devices) at 15 kHz, digitized with a sampling frequency of 50 kHz, and analyzed using pClamp 10.7 software (Axon Instruments).

When a single VDAC channel was reconstituted into PLM and recorded, αSyn was added to both compartments of the experimental chamber to 10 nM final concentration. For data analysis by Clampfit version 10.7, a digital filtering using a 5 kHz low-pass Bessel (8-pole) filter was applied. Individual events of current blockages were discriminated, and kinetic parameters were acquired by fitting logarithmic single exponentials to logarithmically binned histograms as [43]described previously [44]. All lifetime histograms used 10 bins/decade. The number of blockage events for each analyzed current fragment was in the range from 250 to 2500. Three different logarithmic probability fits were generated using different fitting algorithms, and the mean and S.D. of the fitted time constants were used as mean and S.D. for the characteristic open and blockage times. Each channel experiment was repeated at least three times on different membranes. Records for analysis were obtained no less than 30 min after αSyn addition to ensure a steady state. Fits to histograms such as those in Figs. 3C used the maximum likelihood estimator with the simplex algorithm in Clampfit version 10.7.

VDAC ion selectivity was measured in a 1.0 M (*cis*) versus 0.2 M KCl (*trans*) gradient, buffered with 5 mM HEPES at pH 7.4, as described previously [15,45]. The reversal potential – the potential corresponding to the zero-current level – was measured only for single channels under application of either 5 mHz triangular voltage wave of ± 50 mV amplitude or by measuring current acquired at different applied voltages typically at 5 mV intervals. Cl^−^/K^+^ permeability ratio was calculated using the Goldman-Hodgkin-Katz equation as previously described [45]. VDAC voltage-gating was measured on multichannel membranes using a previously described protocol [42,46,47] and in *Supplemental Methods*.

### Thermal fluorescent protein stability assay

The melting temperatures (T_m_) of recombinant VDAC were determined by native tryptophan fluorescence using a Tycho Differential Scanning Fluorometer (Nanotemper). VDAC samples were diluted to 1mg/ml in 25 mM HEPES at pH 7.4, 100 mM NaCl, 0.1% LDAO, 1mM TCEP, and 1mM EDTA buffer. The ratio of native tryptophan 350 nm to 330 nm fluorescence increases at higher temperatures due to exposure of unfolded tryptophan to the solution. The T_m_ was determined from the inflection point of the curve.

### Statistical analysis

Empirical Cumulative Distribution Functions (ECDFs) were calculated based on observed conductances using the Python package iqplot. To determine the confidence interval for the ECDF, 1,000 bootstrap replicates were utilized. The shaded region on the ECDF plots represents the 95% confidence interval derived from the bootstrap analysis. To evaluate the change in substate occurrences for a given mutant or the probability that a channel will display substates for a given number of observations, we modeled the process as being generated by the binomial equation and calculated the maximum likelihood estimation (MLE) for the binomial coefficient. The MLE for Binomial Coefficient to estimate the proportion of channels with substates for each mutant was computed based on 25, 27, and 16 observations for the VDAC2 WT, VDAC2-ΔN, and VDAC2-ΔCys channels, respectively. 10,000 bootstrapped replicates were generated. Significance was determined using a chi-square test.

### Supplemental Material

contains *Supplemental Methods* describing cloning, recombinant protein production, and purification of VDAC2 protein, VDAC reconstitution and conductance measurements, and *Supplemental Figures* S1-S6.

## Results

### VDAC2 dynamically switches between multiple high-conductive open substates

When reconstituted into PLMs formed from a mixture of DOPG:DOPC:DOPE (2:1:1; wt/wt) (2PG/PC/PE) in a symmetrical 1M KCl solution, human recombinant VDAC2 WT forms channels of 3.0 – 4.0 nS conductance which gate, i.e., transition from the high conductance open state to the variety of low conductance “closed” states in response to the applied voltage (Figure 1A, B). This constitutes typical VDAC1 behavior [48,49]. We examined VDAC2 gating in single-channel (Fig. 1A and Supplemental Fig. S1A) and in multichannel membranes (Fig. 1B) to find that the application of 50-60 mV was required to observe typical voltage gating (Fig. 1A and Supplemental Fig. S1A) [49]. The normalized conductance (*G/G_max_*) plots for VDAC2 display the characteristic bell-shaped voltage dependence for all VDACs with enhanced gating, i.e., lower *G_min_*, in response to negative applied voltages (Fig. 1C).

**Figure 1.**
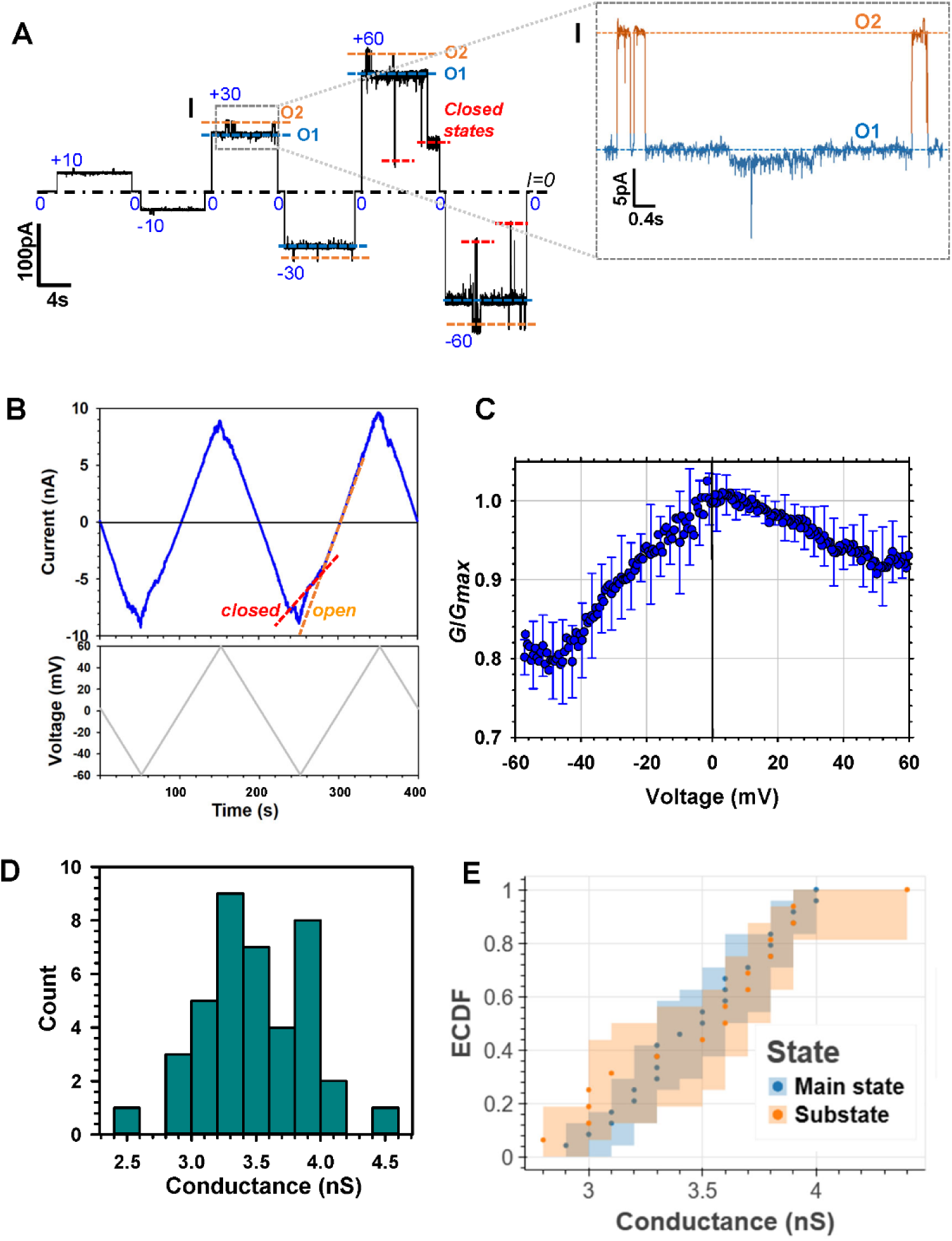
VDAC2 stochastically switches between different “open” conductance substates. **(A**) Representative record of the current through a single VDAC2 WT channel reconstituted into the PLM formed from 2PG/PC/PE lipid mixture in 1M KCl, pH 7.4 at the increasing applied voltages as specified. Two distinct substates of the open state (“O” state) indicated as O1 of 3 ± 0.1 nS (blue dashed line) and O2 of 3.5 ± 0.1 nS (orange dashed line) were observed at all voltages. Characteristic voltage-gating behavior is seen at ±60 mV as stepwise current transition to a low-conductance, “closed” state (red dashed lines). Here and elsewhere the dash-dotted line represents zero current level; dashed lines indicate conductance substates. The current record was digitally filtered at 500 Hz using a low-pass Bessel (8-pole) filter. Inset (**I**): Transitions between two conducting states are shown in a finer timescale using a digital 1 kHz Bessel filter. **(B)** Representative current trace obtained on a multichannel (∼ 40 channels) membrane (upper panel) in response to the applied triangular voltage wave of ±60 mV, 5 mHz (bottom panel) shows a typical VDAC2 WT voltage-gating behavior. Steep slopes at low potentials correspond to the high conductance of open states (orange dashed line) and reduced slopes at higher potential correspond to the lower conductance “closed” states (red dashed line). PLM were made of Polar Lipid Extract (PLE) in 1 M KCl buffered with 5 mM KCl. **(C)** Typical bell-shaped voltage dependence of VDAC2 WT normalized conductance, *G/G_max_*, obtained in multichannel experiments as in (B). *G* is a conductance at given voltage and *G_max_* is the maximum conductance at |*V*| ≤ 10 mV. Data points are means of three experiments ± SD. **(D)** VDAC2 WT single channel conductance histogram. Main (defined as the longest lasting) conductance substates are counted in 24 single channels at different applied voltages. **(E)** The Empirical Cumulative Distribution Function (ECDF) calculated for all main states (blue) and substates (orange) conductances observed in all experiments (N=24 channels). Solid dots denote raw data points. Shaded region represents the 95% confidence interval from 10,000 bootstrap replicates.

The striking difference that we found between VDAC2 and VDAC1 was the appearance of multiple high-conducting open states. They were evident at any applied voltages starting at 10 mV (Supplemental Fig. S1B, C), in approximately 50% of single VDAC2 channels during an observation time of at least 20 min. The example of such spontaneous transitions between two high-conductance states can be seen in inset I in Fig. 1A of the single-channel record of the current through VDAC2. Conductance fluctuates between the long-lasting state O1 of 3.0 nS (indicated by blue dashed lines in Fig. 1, I) and the short-lived state of 3.5 nS (indicated by the orange dashed line). On different single channels (in different PLMs) such substates were observed either for relatively short times, less than 10% of the time spent in the initial state, as shown in the representative trace in Fig. 1A, or for substantially longer times. The existence of multiple substates in VDAC2 results in a relatively wide distribution of open states’ conductances, spanning the range from 3.0 to 4.0 nS (Fig. 1D). Other channels showed a single stable open state throughout the observation time (Figure S1A). To determine if substate conductance distribution depends on the initial state in which the channel inserted in, the Empirical Cumulative Distribution Function (ECDF) plots for the initially observed conductance states (main state, blue) and for the substates (substates, orange) were generated (Fig. 1E). It shows that substate conductance distribution does not depend on the original state in which the channel inserts (main state in Fig. 1E). This suggests that a given VDAC2 channel can initially insert into PLM at any state and then subsequently spontaneously switch to a state of a different conductance.

### VDAC2 high-conductance substates are anion-selective

VDAC1 is known to have multiple voltage-induced “closed” conducting states which are characterized by wide range of conductances and low-anionic or cationic selectivity [46]. This is in a contrast with unique anion-selective open state of VDAC1. To determine whether the observed high-conductance substates of VDAC2 can be reliably attributed to open states and are not voltage-gated states, we examined their selectivity. For this purpose, single channels were reconstituted in a 5x salt gradient (1M KCl *cis* and 0.2 M KCl), where they displayed the same substate behavior as observed in symmetrical salt conditions. A representative trace of VDAC2 obtained in the salt gradient (Fig. 2A) shows two conducting states: the initial long-lasting O1 (indicated by blue dashed lines) and the higher conductance short-lasting state O2 (indicated by orange dashed lines) (inset I in Fig. 2A). The corresponding I/V (current vs voltage) plots for two states O1 and O2 are given in Fig. 2B. The linear regressions determine the slope (conductance) and the reversal potential (*Ψ_R_*, the voltage corresponding to zero current) for each substate. Under these salt gradient conditions, positive *Ψ_R_* indicates anionic selectivity, and negative *Ψ_R_* indicates cationic selectivity. Both conducting substates proved to be anion-selective. Moreover, the lower-conductance substate O1 has a higher reversal potential of 7.2 mV and, accordingly, higher anion selectivity than the higher-conductance substate O2 with *Ψ_R_* = 4.4 mV (Fig. 2B).

**Figure 2.**
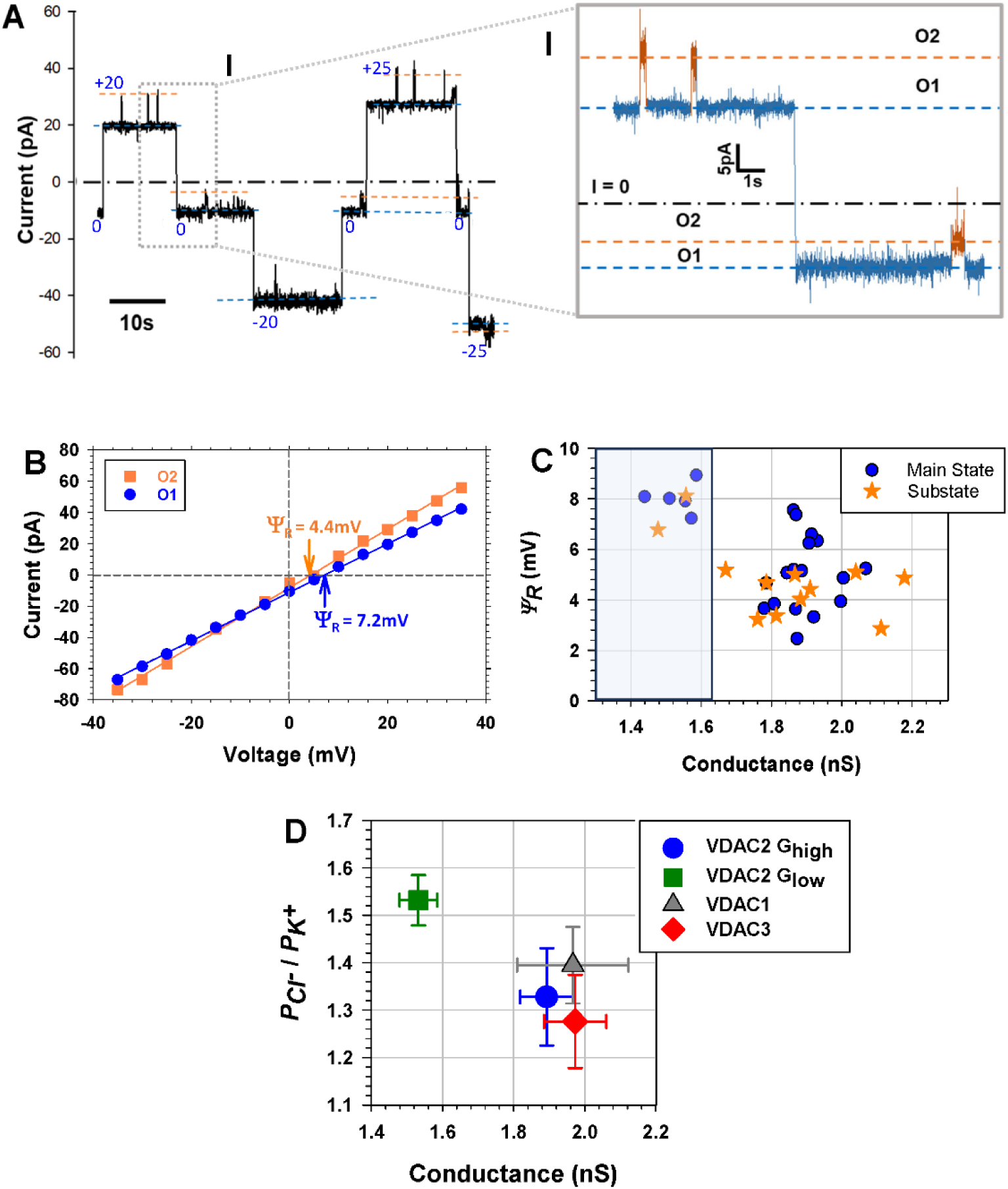
VDAC2 substates are anion-selective open states. **(A)** Representative single-channel trace of the current through VDAC2 WT was obtained using a 1.0 M (cis) / 0.2 M (trans) KCl gradient at different applied voltages, as specified. Blue and orange dashed lines indicate two open substates. Inset (**I**) shows a fragment of ion current (accentuated by a gray box) at a finer time scale, demonstrating two open states, O1 and O2, at +20 mV and 0 mV of the applied voltage. The current record was digitally filtered using a 500 Hz (1 kHz in (I)) low-pass Bessel (8-pole) filter. Other experimental conditions were as in Fig. 1. **(B)**. Current-voltage (*I/V*) curves were obtained for the two states O1 (blue circles) and O2 (orange squares) for the experiment shown in (A). Linear regressions allow for the calculation of the reversal potential (*Ψ_R_*) indicated by arrows for each state. Positive *Ψ_R_* corresponds to anion selectivity. **(C)** Reversal potential (*Ψ_R_*) versus conductance scatter plot for all observed conductance substates: main or long-lasting (blue circles) and short-lasting (orange stars) substates for 19 individual VDAC2 WT channels. (**D**) Comparison of the mean open state conductance and Cl^−^/K^+^ permeability ratio (*P_CL_^−^* / *P _K_^+^*) for each VDAC isoform. The full set of the analyzed VDAC2 conductances was divided into a high conductance state of > 1.6 nS (*G_high_*) and *P_CL_^−^ P _K_^+^* ∼ 1.3 similar to those of VDAC1 and VDAC3, and a state with conductance lower than 1.6 nS (*G_low_*) (highlighted by blue in (C)) and high anion selectivity with *P_CL_^−^ P _K_^+^* ∼ 1.5. Error bars are ± SD from at least 5 independent channel measurements for each VDAC isoform.

The scatter plot of *Ψ_R_* versus conductance for all observed conducting substates obtained in 19 individual VDAC2 WT channels is shown in Figure 2C. *Ψ_R_* is plotted as a function of conductance for the initial or main states (shown as blue circles) that are also typically the most probable states observed for a given channel and for substates (shown as orange stars). The main states and substates could be clustered in at least two groups: one with the higher conductance (>1.6 nS) and lower reversal potential, and another with the lower conductance (<1.6 nS, highlighted in haze blue) and higher *Ψ_R_* (Fig. 2C). It could be seen that substates and main states of the channel occupy the same *Ψ_R_* vs conductance clusters. To empirically determine the validity of such clustering, we performed K-means clustering analysis, with the cluster size determined by the maximum silhouette score. The analysis returned three clusters with the higher conducting states >1.6 nS clustering in two groups and most of the low conducting <1.6 nS states falling into a third cluster (Figure S1D).

These results suggest that the observed substates represent a set of VDAC2 open states characterized by anion selectivity. A comparison of VDAC2 selectivity with VDAC1 and VDAC3 confirms this conclusion. In Fig. 2D it can be seen that the high conductance state of VDAC2 (*G_high_*, which corresponds to typical ∼ 3.8 nS conductance in 1 M KCl symmetrical solution) is similar in conductance and anion selectivity (with Cl^−^ to K^+^ permeability ratio *P_Cl_^−^/P _K_^+^* ∼ 1.4) to the open states of VDAC1 and VDAC3 while VDAC2 lower conductance state (*G_low_*, corresponding to ∼ 3.1 nS in 1 M KCl) is characterized by slightly higher anion selectivity with *P_Cl_^−^/P _K_^+^* ∼ 1.5.

### VDAC2 open substates vary in the kinetics of their interaction with cytosolic regulator αSyn

Previous work demonstrated that while the basic channel properties of VDAC1 and VDAC3 isoforms, such as conductance, ion selectivity, and voltage-gating, are similar, their interaction with VDAC cytosolic protein partners – dimeric tubulin and αSyn – is drastically different, thus reliably distinguishing each isoform [39]. Following these results, we tested the interaction between reconstituted human VDAC2 WT and αSyn at the single-channel level. According to the previously proposed model of αSyn-VDAC complexation [50], the first step of αSyn capture by the VDAC pore is binding of the αSyn N-terminal domain to the membrane surface [33,51] where αSyn preferentially binds to anionic lipids as have been shown in multiple studies [30–32,52,53]. Therefore, to maximize αSyn-VDAC interaction, we used PLM formed from 2PG/PC/PE lipid mixture. The 50% content of anionic DOPG in the lipid mixture increases the on-rate of αSyn-VDAC interaction, allowing for the faster acquisition of statistically reliable data [51]. In addition, another advantage of using PLM of this lipid composition is that VDAC stays mostly open at the applied voltages of up to 60 mV (Fig. 1 and [49]), thus allowing to measure αSyn-VDAC interaction at the wide range of applied voltages. Figure 3A shows a representative trace of VDAC2 WT in the presence of αSyn. Initially, this channel showed two conductance levels, O1 of 3.0 nS and O2 of 3.6 nS, which were recorded at the applied voltages as low as 10 mV. Ten minutes after insertion, the channel spontaneously converted to a state with stable 3.8 nS conductance referred to as O3 (Fig. 3A, Insets I and II). In all three states, the channel is blocked by αSyn, which is manifested by fast current transitions between one of the open states (O1, O2, or O3) and the corresponding blocked states, B1, B2, or B3, of ∼ 0.4 conductance of the corresponding open state (Fig. 1A and *Insets I and II*). Notably, αSyn does not block VDAC2 voltage-induced low-conductance closed state (Supplemental Fig. S2), as was also reported for VDAC1 [29], due to its cationic selectivity. This is another piece of evidence indicating that VDAC2 high-conductance substates are substantially different from the voltage-induced “closed” states of the same single channel.

**Figure 3.**
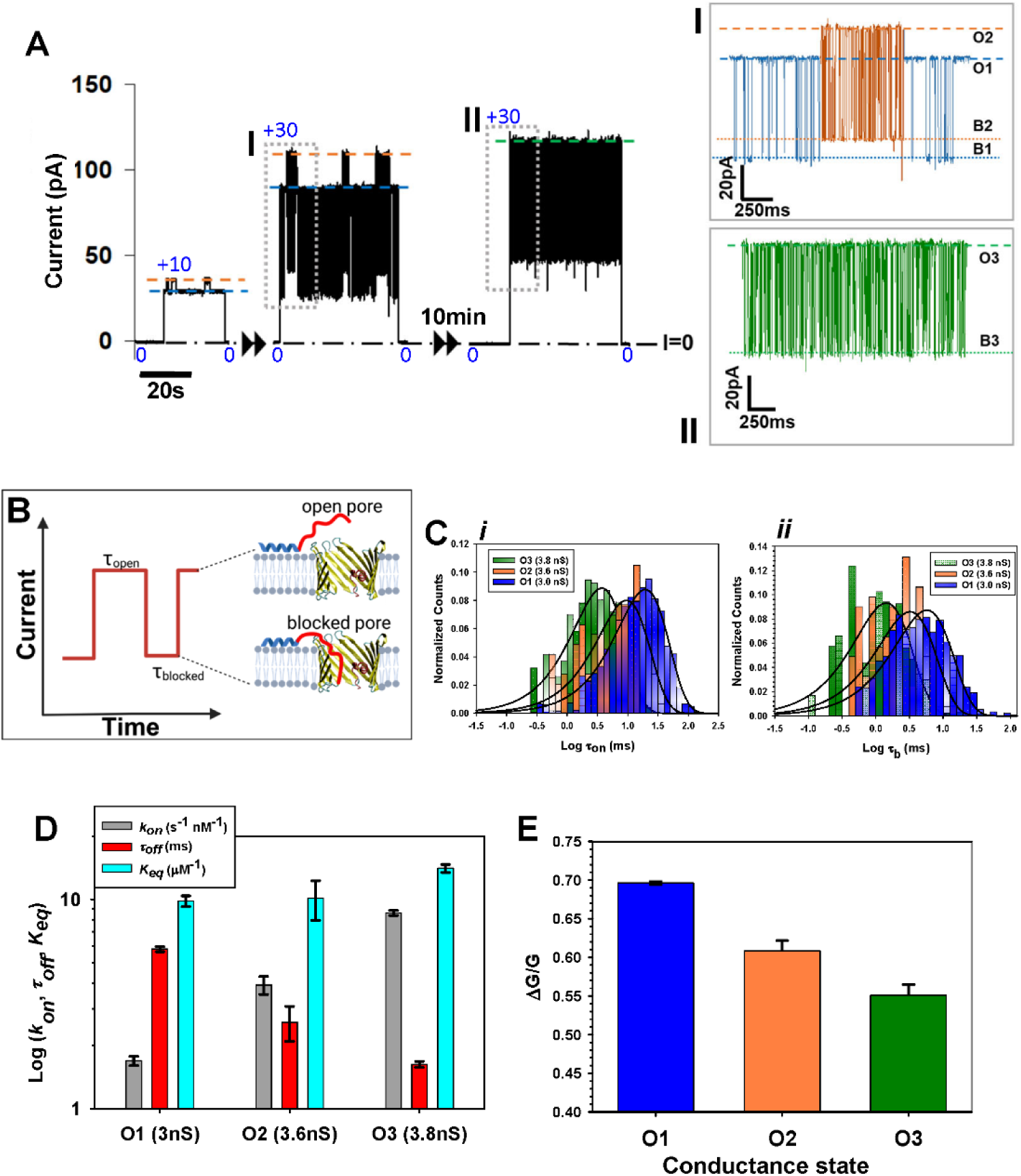
VDAC2 dynamic substates differ from each other in their interaction with cytosolic regulator αSyn. **(A)** Representative current record of a single VDAC2 WT channel, inserted into a 2PG:PC:PE membrane in 1M KCl, pH 7.4 in the presence of 10 nM αSyn added to both sides of the membrane, displays robust substates behavior seen as spontaneous fluctuations between the low-conductance (3nS) long-lasting (O1) and high conductance (3.6 nS) short-lasting (O2) substates at +10 and +30 mV of the applied voltage. After 10 min of recording, the channel spontaneously transitions to a higher conductance (3.8 nS) substate (O3) (trace at the right). Insets **I** and **II** show αSyn-induced blockage events at a finer time scale for each of the three substates at +30 mV (accentuated by gray boxes in the left traces). Each open state (O1, O2, O3) is indicated by a dashed line, whereas respective αSyn blocked states (B1, B2, B3) are indicated by the dotted lines. The current record was digitally filtered using a 500 Hz low-pass Bessel (8-pole) filter. **(B)** A cartoon illustrating that each event of αSyn-VDAC interaction is characterized by the time when the channel is open between each blockage event (*τ_on_*) and the blockage time (*τ_b_*), the duration of each blockage event. **(C)** Representative log-binned histograms of *τ_on_*(i) and *τ_b_* (ii) of each state at +30 mV obtained from the experiment shown in (A). Solid lines are fits to a single-exponential function with characteristic time <*τ_on_*> equal to 19.7, 8.62, and 3.87 ms for state O1, O2, and O3, respectively (i); and *τ_off_* = <*τ_b_*> equal to 5.79, 2.59, and 1.63 ms for state B1, B2, and B3, respectively (ii). (**D**) Comparison of the kinetic parameters: the on-rate, *k_on_* = 1/(<*τ_on_*>[*C*]), where [*C*] is αSyn bulk concentration, the characteristic blockage time, *τ_off_*, and the equilibrium constant, *K_eq_ = k_on_ τ_off_*, of αSyn-VDAC binding for each state at +30 mV. Error bars represent the standard deviation of three separate fitting algorithms for the associated exponential fits. (**E**) Comparison of the αSyn-blocked conductance of each state B1, B2, B3 as shown in panel (A) in I and II. *ΔG/G* is the relative conductance drop, where *ΔG* is the difference between open (*G)* and blocked state conductances of each state O1, O2, and O3. Error bars represent the standard deviation between measurements of *ΔG/G* at more than 3 different voltages.

The important feature of VDAC2 interaction with αSyn is that the frequency of blockage events varies dramatically between the high conductance substates, which can be clearly seen in insets I and II in Fig. 3A with a higher time resolution. Notably, at the same applied voltage, the state with a higher conductance, O3, for a channel shown in Fig. 3A, has a higher frequency of blockage events than the state with a lower conductance, O1. To quantify the αSyn-VDAC interaction, the open times, *τ_on_*, the time between consequent αSyn blockage events, and blockage times, *τ_b_*, (see cartoon in Fig. 3B) are plotted in log-binned histograms for each state at the applied voltage of +30 mV (Fig. 3C). State O1 has the longest intervals between αSyn blockage events, followed by O2 and finally O3, demonstrating that *τ_on_* decreases with the state’s conductance increase (Fig. 3C, i). The blockage times follow the opposite sequence: state O1 has the longest *τ_off_* = < *τ_b_* > followed by O2 and O3 (Fig. 3C, ii). The comparison of the kinetic parameters of αSyn-VDAC binding for each state obtained in the experiment in Fig. 3A is shown in Fig. 3D. The on-rate constant of αSyn capture by the pore, *k_on_* = 1/(<*τ_on_*>[*C*]), where [*C*] is αSyn bulk concentration, and the characteristic off-rate, *τ_off_*, were calculated by averaging the rates determined via three separate fitting algorithms for the log-probability distribution (Fig. 3C) (see Methods). As conductance increases from state O1 to O2 to O3, *k_on_* increases from 1.70 to 3.90 to 8.62 s^−1^ nM^−1^, respectively, and *τ_off_* decreases from 5.79 ms in O1 to 2.59 ms in O2 to 1.63 ms in O3. Importantly, the equilibrium constant, *K_eq_* = *k_on_ τ_off_* (light blue bars in Fig. 3D), varies negligibly between the substates. To compare the trend in *k_on_* values across multiple substates in individual VDAC2 channels, *k_on_* values were plotted versus corresponding conductance (Supplemental Fig. S3A). In every case, (7 independent experiments with different channels, and total 12 analyzed substates), as substate conductance increased, *k_on_* also increased with 95% confidence (Supplemental Fig. S3C), confirming the results obtained on the same channel (Figure 3D). This trend is consistent across multiple observed single-channel measurements despite large variability in absolute values of the kinetic parameters between individual channels and, correspondingly, a low coefficient of determination R^2^ = 0.33.

For VDAC1 and VDAC3, the conductance of the αSyn blocked state is ∼0.4 of the open state conductance [29,39], giving *ΔG/G* ∼ 0.6. A comparison of the blocked state’s relative conductance, *ΔG/G,* for each state O1, O2, and O3 (as in Fig. 3A), is shown in Fig. 3E. State O1 has the highest *ΔG/G* with 0.70 of conductance blocked, followed by O2 with 0.61, and O3 with 0.55. A comparison of *ΔG/G* values across different substates using the same set of individual VDAC2 channels as for *k_on_* and *τ_off_* analysis confirms the trend found for the same channel (Supplemental Fig. S3D). Statistical analysis of the substates was performed using the nonparametric paired Wilcoxon signed rank test, comparing substates observed on the same channel, allowing comparison between data taken at different voltages (Supplemental Fig. S3A-B).

VDAC2 has a lower average *k*_on_ of aSyn blockage than VDAC1 but a larger variance (Supplemental Fig. S4A). At negative applied voltages, the ratio between the highest *k*_on_ and the lowest one is up to ∼3 times. At positive applied voltages, there is an even larger variability between channels, namely, up to ∼10 times difference in *k*_on_ between different channels, reflecting the intrinsic asymmetry of VDAC channel [42]. Nevertheless, the range of *k*_on_ values for VDAC2 is below those for VDAC1 obtained under the same conditions [51].

### VDAC2 N-terminal extension and extra cysteines affect the occurrence of conductance substates but do not eliminate them

VDAC2 differs most prominently from VDAC1 by the presence of an extra 11-amino acid NTE and seven extra cysteine residues, six of which face intermembrane space in mitochondria [10,38] (Fig. 4A). These features have been proposed to contribute to VDAC2’s unique function [11,54] and have already been shown to alter the chemical-physical properties of the hVDAC2. Maurya et al showed that cysteine-less hVDAC2 protein has higher thermostability and, consequently, a more stable β-barrel than the WT [55]. They also showed that the 11-residue NTE plays a role in hVDAC2 refolding in micelles and lipids. However, neither cysteines nor NTE significantly affect VDAC2 basic properties such as single-channel conductance or voltage-gating. Based on these data, we decided to determine whether these sequence features are responsible for VDAC2’s dynamic substate behavior. We utilized two previously characterized constructs: hVDAC2 with truncated ΔN-terminus 1-11 (ΔN-VDAC2) and a mutant lacking all cysteines (VDAC2-ΔC). Both ΔN-VDAC2 and VDAC2-ΔC mutants reconstituted into the same 2PG/PC/PE PLM display channels of 3.8-4.0 nS conductance. However, lower conductance states were still observed, prevalently in the ΔN-VDAC2 mutant and less so in the VDAC2-ΔCys mutant.

**Figure 4.**
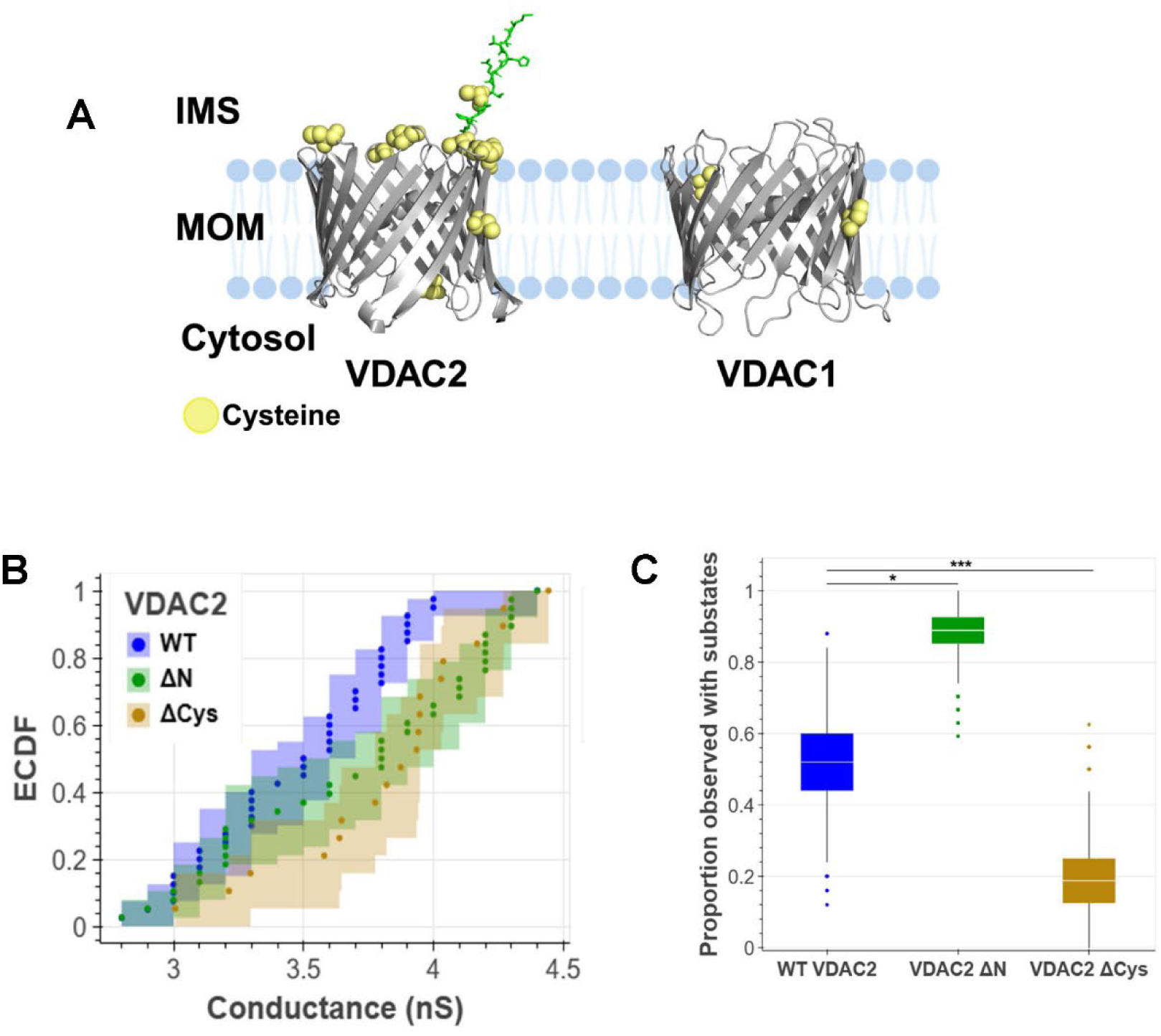
N-terminal extension and cysteine mutations influence but do not abolish VDAC2 substates. **(A)** Structure of human VDAC2 (Alphafold2 prediction) and VDAC1 (hVDAC1: PDB ID: 2JK4). Cysteines are shown as yellow bolls. NTE of VDAC2 is shown in green. **(B**) ECDF was calculated using conductance values of the measured substates for all conductances observed for WT (blue), ΔN (green), and ΔCys (goldenrod) VDAC2 channels. Occurrence was liberally defined as whether a state transition had been observed during the entire recording period for a given channel. Out of 24 single-channel recordings of VDAC2 WT, 13 channels displayed substate behavior, compared to 25 channels out of a total of 27 for ΔN-VDAC2 channels and 3 out of 16 channels of VDAC2-ΔC. Solid dots represent raw data points. The shaded region represents the 95% confidence interval from 1000 bootstrap replicates. **(C)** Box and Whisker plots for MLE probability of the binomial coefficient; substate behavior being observed in WT (blue), ΔN (green), and ΔCys (goldenrod) VDAC2 channels. 10,000 bootstrapped replicates were simulated from a maximum likelihood estimation of the binomial coefficient originally calculated from 25, 27, and 16 observations of WT, ΔN, and ΔCys VDAC2, respectively. The top and bottom of the box are, respectively, the 75^th^ and 25^th^ percentiles of the data. The line in the middle of the box is the median. The top whisker extends to the maximum of the set of data points that are less than 1.5 times the interquartile regions beyond the top of the box, with an analogous definition for the lower whisker. Data points not between the ends of the whiskers are plotted as individual points. Substate occurrence between mutants was compared with a chi-squared test. “*” indicates p-value < 0.05, “***” indicates a p-value < 0.001.

To compare substate occurrence between two mutants and VDAC2 WT, the ECDF for two mutants and the WT VDAC2 was calculated (Fig. 4B). This allows to directly compare proportions of channels observed with substates. The analysis shows that ∼93% of ΔN-VDAC2 channels have substates in comparison with ∼54% of VDAC2 WT and ∼19% of VDAC2-ΔC channels (Fig. 4C). Thus, the removal of the first 11-amino acids greatly enhances substate occurrence. In contrast, the removal of cysteines reduces, though less significantly, the occurrence of substates. Neither mutation abolishes the occurrence of substates, and, therefore, these structural features of VDAC2 are not the source of substate appearance. However, the NTE and cysteines can allosterically regulate the dynamics of this behavior. The ΔN-VDAC2 channel is less thermostable than VDAC2 WT [11], while the VDAC2-ΔC mutant had been shown to have a higher melting temperature (T_m_) [13,55], thus suggesting that the presence of NTE and extra cysteines might influence the occurrence of the substates independently of their effect on β-barrel thermostability.

### N-terminal extension and cysteines affect the VDAC2 interaction with αSyn

We showed previously that the removal of cysteines from hVDAC3 altered that channel’s interaction with αSyn [39]. The on-rate of αSyn interaction with the VDAC3 cysteine-less mutant was ∼ 10x higher than for the WT, but only at negative potentials [39]. Therefore, the next logical question was whether cysteines and NTE could affect VDAC2’s interaction with αSyn. Both ΔN-VDAC2 and VDAC2-ΔC mutants exhibited characteristic reversible blockages in the presence of αSyn (Fig. 5A, C, D). ΔN-VDAC2 displayed long-lasting substates with a visible difference in interaction kinetics with αSyn (Fig. 5A and Inset I). The representative trace of a single ΔN-VDAC2 channel in Fig. 5A shows three substates, O1, O2, and O3, at the applied voltages of ±10 and ±35 mV. The ΔN-VDAC2 mutant exhibited two predominant long-lasting substates of 3.2 and 4.0 nS in different channels, allowing the measurements of the kinetic interaction with αSyn for both substates at the wide range of applied voltages on different single channels (Fig.5B). As observed for VDAC2 WT (Fig. 3D), the higher conductance substate of 4.0 nS has higher *k_on_* than the lower conductance substate of 3.2 nS at all voltages of both polarities (Fig. 5B, i).

**Figure 5.**
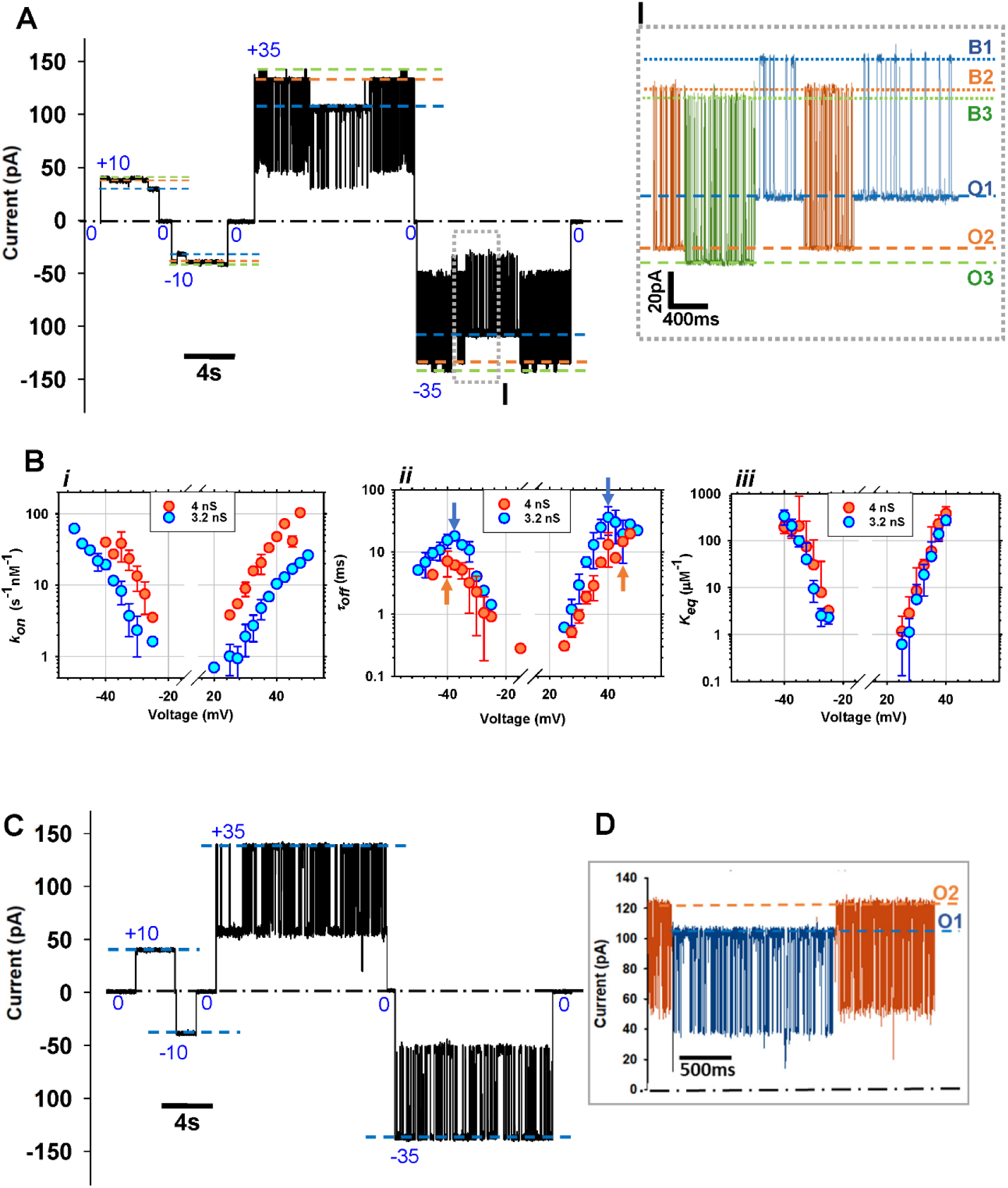
αSyn blocks the higher-conductance substates with a higher on-rate than the lower-conductance states across VDAC2 mutants. **(A)** Representative single-channel current trace obtained with the reconstituted ΔN-VDAC2 mutant with 10 nM αSyn added to both sides of the membrane. Three distinct substates, O1, O2, and O3 are observed at ±10 and ±35 mV of the applied voltage. Inset (**I**) shows αSyn-induced blockage events at a finer time scale for each of the three observed substates at - 35 mV (accentuated by the gray box in the left trace). Open states O1, O2, and O3 can be observed along with their corresponding αSyn blocked states B1, B2, and B3. Current record was digitally filtered using a 1 kHz low-pass Bessel (8-pole) filter. (**B**) Voltage dependences of the on-rate constant *k_on_* (i), the mean blockage time *τ_off_* (ii), and the equilibrium constant *K_eq_* (iii) (calculated for the retraction regime only) of αSyn binding to two ΔN-VDAC2 substates with conductances of 4 and 3.2 nS. Arrows in (ii) indicate the transitions between blockage/retraction and translocation regimes. Data are means of three independent experiments ± SD. **(C)** A representative current record of VDAC2-ΔC single channel in the presence of 10 nM αSyn on both sides of the membrane showing one open state of 4 nS. (**D**) Example of the current record of VDAC2-ΔC with two substates of 3.5 and 3 nS obtained at +35 mV of the applied voltage. Current records were digitally filtered using 500 Hz (C) and 1 kHz (D) low-pass Bessel (8-pole) filters.

The mean blockage time *τ_off_* for VDAC2 mutants and the WT is also highly voltage-dependent and has a characteristic biphasic behavior (Fig. 5B, ii and Supplemental Fig. S4B) as it was shown for VDAC1 and VDAC3 [29,39]. At low applied voltages, *τ_off_* increases with voltage, which corresponds to the blockage/retraction regime when αSyn molecule is captured by the VDAC pore and then released back to the same side of the membrane [33,34,56]. At higher applied voltages – usually |*V*|> 40 mV depending on the VDAC type and experimental conditions – *τ_off_* decreases with voltage, corresponding to the translocation regime. This happens because under the high applied voltage, the N-terminus of αSyn detaches from the membrane, allowing the whole αSyn molecule to translocate to the opposite side of the membrane [34]. The 4 nS substate of ΔN-VDAC2 has a shorter *τ_off_* than the 3.2 nS substate at all applied voltages (Fig. 5B, ii). This results in the unchanged equilibrium constant *K_eq_*, calculated for the retraction regime only, which varies insignificantly between two substates across applied voltages (Fig 5B, iii), similar to the substates analyzed on the same VDAC2 WT channel (Figure 3D). These results indicate that while substate transitions can reliably affect the kinetics of the αSyn-VDAC2 interaction, the final energy change, or affinity between αSyn and the VDAC2 pore remains unaltered.

The VDAC2-ΔC mutant also exhibited conductance substates, but less frequently than ΔN-VDAC2 and the WT (Fig. 4C). An example of two substates in VDAC2-ΔC channel of 3.0 and 3.5 nS is shown in Fig. 5D. The on-rate of αSyn blockages was again higher in the higher-conductance substate than in the lower-conductance one. Notably, the conductance of this predominant state is typical for the open-state conductance of VDAC1.

### E84A mutation increases β-barrel thermostability of VDAC2 but does not abolish the substates

One of the differences between mammalian VDAC isoforms is the charged glutamate in position 73 for VDAC1 and position 84 in VDAC2 and a non-charged glutamine in position 73 in VDAC3 [57]. The E84 residue in VDAC2 corresponds to the notorious E73 of VDAC1 – a charged residue buried in the middle of the hydrophobic membrane and continuously under investigation to determine its physiological relevance [49,58–60]. E73 was implicated in the stability of the VDAC1 β-barrel, whose dimeric channel interactions appear to be largely determined by this residue [61,62]. Previous work on VDAC gating has demonstrated that β-barrel dynamics are often the result of subtle allosteric network interactions across the channel [63]. E84 in VDAC2 is thought to be involved in biologically relevant interactions such as those with ceramide (Dadsena, Bockelmann et al. 2019). We hypothesized that if conductance substates are due to the dynamic rearrangement of the salt bridge ensemble inside the pore leading to the set of pores of different selectivities and conductances, it is natural to expect a significant involvement of the E84 residue. Following this hypothesis, we tested the possible involvement of this residue in conductance substates formation in VDAC2 by using VDAC2 mutant E84A which replaces the membrane facing glutamate. E84A displayed enhanced thermostability, as measured by native tryptophan scanning fluorimetry (Fig. 6A). Its thermostability is significantly higher than that of VDAC1 and VDAC2, indicating the reduced flexibility of the β-barrel. However, substates were still observed (Fig. 6B and Supplemental Fig. S5B). The VDAC2 E84A mutant gates similarly to the WT in single and multichannel experiments (Supplemental Fig. S5B). Thus, the E84 residue does not affect VDAC2 voltage gating, consistent with observations in the E73A VDAC1 mutant voltage gating experiments [49]. The VDAC2 E84A mutant displayed more stable and long-lasting substates than the WT, similarly to the substate stabilization observed in ΔN-VDAC2. An example of the E84A mutant displaying two long-lasting stable substates of 3.0 and 3.6 nS is shown in Fig. 6B in the presence of 10 nM αSyn. Similar to VDAC2 WT and the ΔN-VDAC2 and VDAC2-ΔC mutants, the substate with higher conductance of 3.6 nS in the E84A mutant has higher *k_on_* and shorter *τ_off_* than the lower conductance substates of 3 nS, resulting in unchanged *K_eq_* (Fig. 6C). These findings indicate that similar to NTE and extra cysteines, E84 mutation affects β-barrel flexibility but does not eliminate substate appearance.

**Figure 6.**
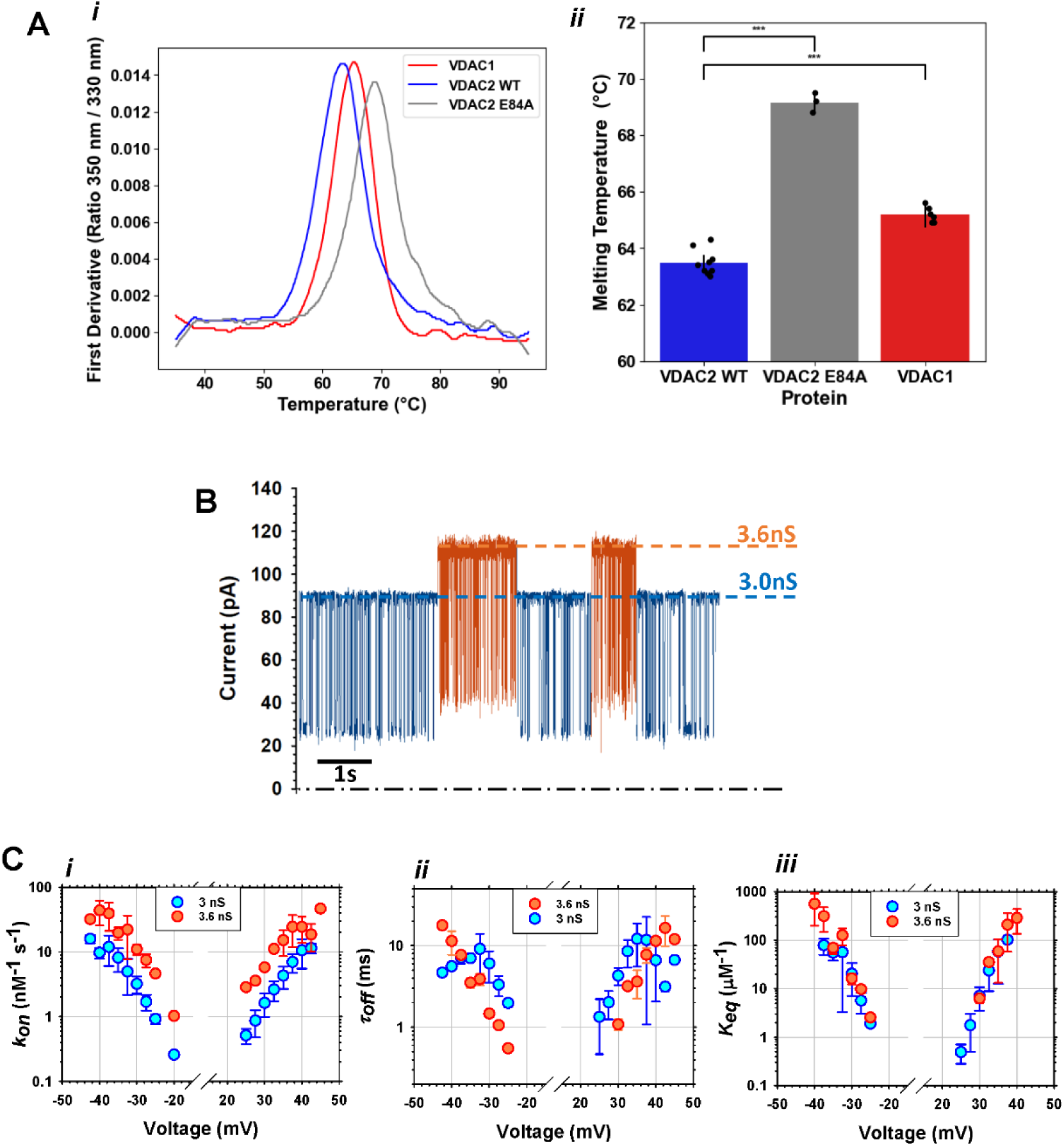
E84 residue of VDAC2 contributes to β-barrel stability but does not affect substates appearance. (A) **(i)** The first derivative of the ratio of 350 nm / 330 nm tryptophan fluorescence vs temperature demonstrates the inflection melting point (*T_m_*) for VDAC1 (red), VDAC2 WT (blue) and VDAC2 E84A (gray). (**ii**) Box and scatter plot demonstrating average *Tm* for VDAC1 (red), VDAC2 WT (blue), and VDAC2 E84A (gray). Melting temperatures were compared with Student’s T-Test; ‘***” indicates a p-value < 0.000001. **(B)** The representative current record of the VDAC2 E84A mutant showing two equally long-lasting conducting states of 3.0 and 3.6 nS in the presence of 10 nM αSyn on both sides of the membrane at the applied voltage of 30 mV. The current record was digitally filtered using a 1 kHz low-pass Bessel (8-pole) filter. (**C**) Voltage dependences of the on-rate constant *k_on_* (i), the mean blockage time *τ_off_* (ii), and the equilibrium constant *K_eq_* (iii) of αSyn binding to two VDAC2 E84A states with conductances of 3.6 and 3.0 nS. Data are means of three independent experiments ± SD.

## Discussion

VDACs have been implicated in diverse mitochondrial signaling pathways, with VDAC2 being crucially important in embryonic development [6,9]. Yet, the biophysical underpinnings of its special role in regulating functions remain unexplored. In this study, we revisited the basic biophysical properties of recombinant human VDAC2 using single-molecule electrophysiology to demonstrate the unique plasticity of this channel’s properties and its interaction with cytosolic proteins. Our data show that VDAC2 stochastically switches between multiple high-conductance states, which results in a wide conductance distribution of the “open” state uncommon for other VDACs. The existence of VDAC2 multiple open states (but not the spontaneous transitions between them) was reported previously [12], with an average conductance of ∼ 3.5 nS in 1 M KCl at room temperature, which is lower than the typical open state conductance of VDAC1 and VDAC3 at these conditions. We verified that these states are not the conventional VDAC’s voltage-gated states because they are even more anion selective than typical ∼4 nS open states of VDAC1 or VDAC3 (Fig. 2C) [39] as opposed to the more cationic voltage-gated states. Besides, they are observed at all applied voltages starting from as low as 10 mV (Fig. 1, 2). Under sufficiently high applied voltage, VDAC2 characteristically gates similarly to its other two isoforms [39,49] and to VDACs isolated from their native mitochondria of different species - fungus, yeast, or plant [46,64–66]. This is manifested by a bell-shaped conductance vs voltage (*G/V*) dependence (Fig. 1C) and single-channel current transitions from the open state to the variety of closed states under the applied voltages of |*V*| ≥ 60 mV (Fig. 1A). Therefore, the distinctiveness of VDAC2 among the three isoforms is that instead of the unique high-conductance anion-selective state, VDAC2 possesses a variety of such states. Notably, the characteristic duration and probability of each conductance state varies significantly between individual channels. The selectivity of the states with high conductances, which appeared to be in the range of open state conductances of VDAC1 and VDAC3 (1.8 – 2.1 nS in 1.0 M /0.2 M KCl salt gradient), fits into the range of VDAC1 and VDAC3 selectivity with permeability ratio *P_Cl-_ /P_K+_* =1.35 ± 0.15, while the VDAC2 substates with lower conductance of 1.5 ± 0.05 nS have, on average, even higher anionic selectivity with *P_Cl-_ /P_K+_* = 1.5 ± 0.05 (Fig. 2 D).

Using αSyn, a known potent cytosolic regulator of VDAC1 and VDAC3, as a sensitive molecular probe of the VDAC pore, we explored different substates of VDAC2. We found that αSyn characteristically blocks all registered high-conductance VDAC2 substates. Notably, αSyn does not block conventional voltage-gated low-conductance states either in VDAC2 or other VDACs [29]. This, together with anionic selectivity and high conductance, proves that these VDAC2 substates differ from the voltage-induced, more cation-selective lower-conductance states. Our results also show that αSyn quantitatively discriminates between different high conductance states: the on-rate of αSyn capture by the VDAC2 pore rises with the substate conductance while the blockage time decreases (Fig. 3D). This correlation is especially clear if the on- and off-rates for different substates are compared on the same VDAC2 WT channel. Across multiple individual channels of VDAC2, considerable variability in kinetic parameters exists, especially in *k_on_* values, suggesting that channel structure is highly dynamic.

Considering that VDAC permeability to mitochondrial metabolites, such as ATP, ADP, and Ca^2+^, depends on VDAC pore anion/cation selectivity, we can speculate that by switching between substates, VDAC2 is capable of fine-tuning the regulation of metabolite and Ca^2+^ transport in and out of mitochondria [40]. If such a physiologically meaningful mechanism exists, the immediate questions to answer concern the structural features of the substates’ appearance and the physiological factors that regulate these substates in VDAC2 in live cells.

### Possible conformational changes in VDAC2 leading to substate appearance

In an attempt to answer the first question, we studied the effect of the apparent sequence differences between VDAC2 and VDAC1 – the 11-amino acid NTE and the seven extra cysteines – on the formation of substates. We found that both NTE and cysteines quantitatively affect substates’ appearance frequencies (Fig. 4) with their substantial decrease in VDAC2-ΔC down to ∼19% and increase up to ∼92% in ΔN-VDAC2, compared with ∼54% of substates occurrence for the WT. The absence or presence of NTE and extra cysteines influences substates’ appearance but does not eliminate them, thus suggesting that neither is the structural reason for substates’ formation. Furthermore, the deletion of NTE, in fact, stabilizes two predominant conductance substates in different ΔN-VDAC2 channels.

The VDAC2 E84A mutant is more thermostable than the VDAC2 WT and VDAC1 (Fig. 6A). Still, the reduced flexibility of its β-barrel does not eliminate conductance substates. Like in ΔN-VDAC2, E84A mutation results in the stabilization of two predominant substates of 3 and 3.6 nS. The kinetic analysis of αSyn interaction with ΔN-VDAC2 and E84A mutants confirms the results obtained in VDAC2 WT. They also exhibit a positive correlation between conductance and *kon* and a negative correlation between conductance and *τ_off_*, resulting in the same *K_eq_* for both conductance levels. The results show that VDAC2’s NTE, cysteines, and E84 residue all contribute to, but are insufficient, to determine VDAC2’s dynamic behavior. Based on the available structural and computational studies on VDAC1 [61–63,67,68] and the predicted structural similarity between VDAC1 and VDAC2 [38] shown experimentally for VDAC2 zebrafish [69], we can only speculate that the variety of conductance substates in VDAC2 may arise from dynamic rearrangements in the network of charged residues and salt bridges inside the pore in the proximity to the restriction zone and the tentative selectivity filter formed by two N-terminus α-helices as in VDAC1 [63,70–72]. Importantly, the conductance substates were also observed for zebrafish VDAC2 (Supplemental Fig. S6).

Furthermore, αSyn recognizes these substates by interacting from both sides of the bilayer, implying that the structural changes underlying the states’ appearance most likely occur close to the center of the channel rather than in the asymmetric loops on either cytosolic or mitochondrial-facing sides. The long-awaited atomic structure of mammalian VDAC2 will help to test these speculations.

One of the important results is that despite the significantly different on-rates of αSyn capture by different VDAC2 substates, the equilibrium constants *K_eq_* remain virtually unaltered. This can be explained by the transition path energy landscapes shown in Fig. 7 for two VDAC2 substates. *K_eq_* is an exponential function of the change in the free energy *ΔΔUU* between the minima for the membrane-bound “free” αSyn molecule (potential well on the left) and for the pore-trapped one (potential well on the right). Because the experiment does not show any differences in the equilibrium constant in different substates, this means that *ΔΔUU* is conserved, suggesting that the physical state of the αSyn molecule trapped within the VDAC pore is the same for all the substates. However, the profound changes in the on- and off-rates indicate that the region of the highest energy (transition state) that the synuclein molecules must cross before reaching their final state in the pore differs for different substates. We can estimate the change in the transition state energy for substates 1 and 2, ΔΔ*U* ^(1,2)^, from *k_on_* values as

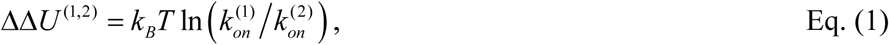

Where 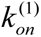 and 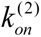 are the on-rates in substate (1) and (2), respectively. For example, applying Eq. (1) to the WT channel data presented in Fig. 3, we find that the change in the transition state energy between the open substate of the highest conductance, O3, and that of the lowest conductance, O1, is 1.6 *k_B_T*.

**Figure 7.**
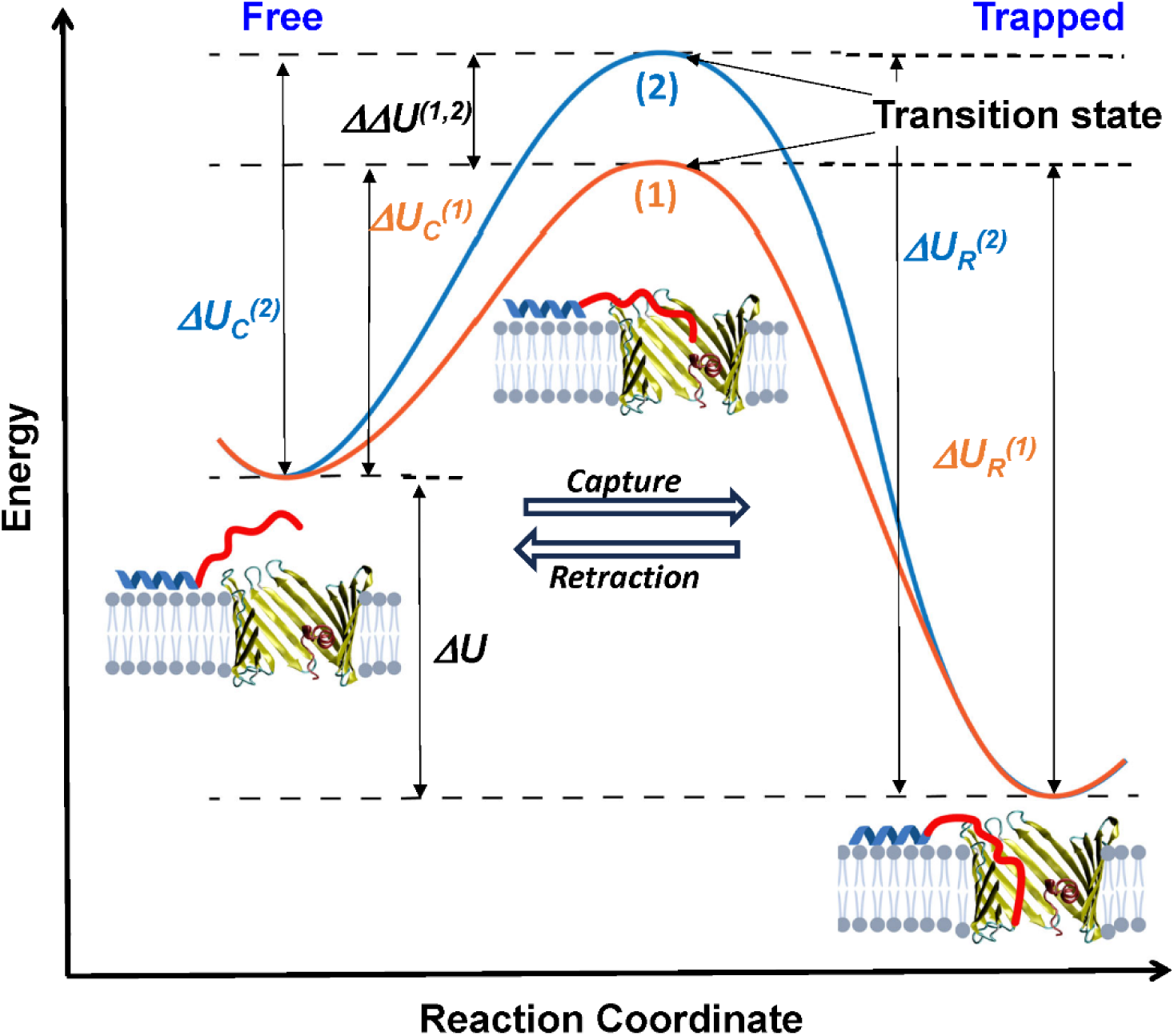
Changes in the transition state energy barrier underlie the kinetic differences in the αSyn-VDAC interaction between substates. Free energy landscape diagram representing a model of the αSyn-VDAC interaction as a function of the reaction coordinate. The potential well of the initial state of free αSyn bound to the membrane is shown on the left. The potential well of the final state, whereupon the αSyn molecule is captured by the VDAC pore, is shown on the right. The energy landscape experienced by the αSyn molecule as it complexes with the higher conductance VDAC2 substate (1) is represented in orange, whereas the landscape for the lower conductance substate (2) is represented in blue.

On the one hand, empirical observations on αSyn-VDAC interaction [29,51] show that the on-rate is a strong exponential function of the applied voltage. On the other hand, in the case of open VDAC2 substates studied here, the on-rate is very sensitive to the ionic current through the channel, even at the same applied voltage. Specifically, it increases up to an order of magnitude for the higher currents, corresponding to the substates of higher conductance. In both scenarios – increasing applied voltage and increasing current at the same voltage – the fraction of the applied voltage that projects into the bulk solution in the vicinity of channel entrances increases. This is especially important in the case of VDAC because of the quite substantial access resistance of this channel, as was shown in experiments with water-soluble polymers that change the specific conductivity of the bulk solution [73,74] and confirmed within a mean-field approach by solving three-dimensional Poisson and Nernst-Planck equations [74]. The fraction of the applied potential extending into the bulk changes the local concentration of charged molecules in the vicinity of the channel entrance and thus is able to change their capture rates. A natural question is whether the field at the channel entrance is strong enough to account for the observed effects and contribute significantly to the free energy profile in Fig. 7.

Let us first analyze the on-rate voltage dependence presented in Figs. 5B and 6C. The total channel resistance, *R*, is the sum of access resistances at both entrances of the channel, 2*R_acc_*, and the resistance of the channel proper, 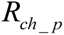 [75]

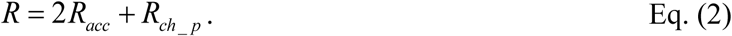

Both experimental work and the mean-field calculations performed for VDAC in 1 M KCl give the value of 2*R_acc_* as 19 to 20 % of the total channel resistance, that is 2*R_acc_* ; 0.2*R*, which means that the access resistance at each channel entrance is close to 0.1 *R*. Therefore, ∼10% of the applied voltage drops at the channel entrance. The data in Figs. 5B and 6C show that upon a 10 mV increase in the (modulus of) applied voltage, the *k_on_* increases by a factor of ∼8. Using Eq. (1) for ΔΔ*U* ^(1,2)^, where indices (1) and (2) now refer to the heights of the barriers for αSyn capture, Δ*U_C_*, at the two voltages differing by 10 mV, we arrive to 1.7 *k_B_T*. If we now assign the on-rate increase to the access resistance effect, then recalling that the change in the fraction of potential extending from the channel is 10% only, that is, *V_acc_* ; 1 mV, and using

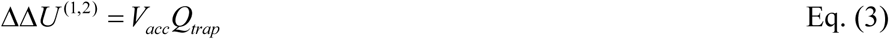

we arrive at ∼ 50 elementary charges for the effective “trapping charge”, *Q_trap_*. This charge significantly exceeds the total charge of the αSyn C-terminus, meaning that the applied voltage effect on the height of the transition state in Fig. 7 is more complex.

Let us now look at the change in the on-rates between the open substates using the same line of reasoning. As explained above, at 30 mV of applied potential the transition of the VDAC2 channel from the open substate O1 of conductance *G*^(*O*1)^ = 3.0 nS to substate O3 of conductance *G*^(*O*3)^ = 3.8 nS decreases the barrier by 1.6 *k_B_T*. The change in the *V_acc_* can be estimated through the change in the channel current, Δ*I* = _(_*G*^03^ − *G*^01^ _)_*V* as *V_acc_* = *R_acc_*Δ*I* = 0.65 mV. Using Eq. (3), we arrive at *Q_trap_* of ∼ 60 elementary charges. Again, this number is too large for the access resistance effects to explain the on-rate change between the states. However, the closeness of the two charge estimates, namely 50 vs 60, strongly suggests a similar physical mechanism of on-rate regulation in both cases. The capture is governed by the voltage distribution along the channel and within its access areas, which defines the physics of the observed phenomena. The exponential dependence of the on-rate on voltage and a factor of four overestimation of *Q_trap_*, compared to the actual charge of the αSyn C-terminus, means that the capture of a disordered protein molecule by the channel is a complex process dominated by the entropic barrier [76] and that the transition state in Fig. 7 corresponds to the C-terminus being partially captured by the pore. This way, the potential drop over the terminus is much larger than the potential drop over the VDAC access resistance.

The significant differences in Δ*G* / *G* found for different substates (Fig. 3D) also suggest that the substates should vary by structural rearrangements inside the pore. Indeed, one may expect that the narrower, less conducting channel will be influenced by the trapped polypeptide to a greater degree than the wider, more conducting one.

It is worth noting here that a similar correlation between the channel conductance and the on-rate of αSyn molecule capture was previously demonstrated for three different β-barrels formed by α-Hemolysin, MspA, and VDAC [50]. At the applied voltage of 35 mV, VDAC1, the channel of the highest conductance of 4 nS in 1M KCl has about three orders of magnitude higher on-rate than that for α-Hemolysin, the channel of the lowest conductance of 0.73 nS in the same salt. However, remarkably, the equilibrium constant was not conserved for these three channels, opposite to what is reported here for the VDAC2 substates. We conclude that the regulation of all substates of the open conformation of VDAC2 by αSyn is described by the same affinity, suggesting that once the αSyn molecule is captured, its physical state and interactions in the pore are the same for all substates, but differs significantly by its kinetic parameters.

### Physiological implications of VDAC2 unique plasticity

The unexpected finding that substates vary by the kinetics of αSyn’s interaction with VDAC2 but not its affinity implies that substates do not tune αSyn regulation of channel fluxes. However, the altered kinetics demonstrate that the capture of the synuclein molecule is indeed affected by the structural changes in the channel. It is natural to expect that such conformational changes in VDAC2 will affect interactions of other proteins with the channel. Potential interactors may include pro-apoptotic proteins BAK and BAX, which could selectively recognize and bind a particular substate of VDAC2. Experimental validation of the BAX/BAK interaction at the single molecule level remains challenging due to the pore-forming nature of these molecules. Nevertheless, the dynamic conformational flexibility identified in the present study may allow VDAC2 to recognize many such binding partners. It would be of great importance to identify VDAC2-specific interactors using proximity labeling or immune precipitation mass spectrometry experiments.

The question as to why VDAC2 knockout is lethal or partially lethal, but knockouts of the other two isoforms are not, remains open. We speculate that VDAC2’s unique plasticity, manifested by spontaneous transitions between high conductance substates revealed in in vitro single-channel experiments, could be a key to understanding its physiological significance. These data could tentatively explain the unique functional role of VDAC2 through its ability to dynamically adapt to rapidly changing metabolic cell conditions and to alter the rate of interaction with its presumably multiple protein partners.

## Author contributions

W. M. R, T. K. R., and S. M. B. designed the experiments; W. M. R. performed the experiments; M. G. L. performed experiments with zfVDAC2; W. M. R. and T. K. R. analyzed the data; W. M. R., T. K. R., and S. M. B. wrote the manuscript; R. M., H. K., and T.-Y. D. Y. produced VDAC proteins; all authors contributed intellectually, revised, and edited manuscript.

## Acknowledgments

W.M.R, M.G.L, T.K.R. and S.M.B. were supported by the Intramural Research Program of the *Eunice Kennedy Shriver* National Institute of Child Health and Human Development of the National Institutes of Health. This research was funded in part by a generous gift from the J. Yang and Family Foundation.

## Notes

### Competing Interest Statement

The authors have declared no competing interest.

