## Supplemental Material for "Conformational plasticity of mitochondrial VDAC2 controls the kinetics of its interaction with cytosolic proteins"

**Supplemental Methods**

**Purification of VDAC2 mutants**

Cells were resuspended in lysis buffer (50 mM Tris-HCl, 250 mM NaCl, 10 mM 2-mercaptoethanol (Bioshop, Burlington), protease inhibitors tablets (Roche, Basel), pH 7.0 and lysed by a French press (Genizer, Irvine). The VDAC2 containing inclusion bodies were isolated using sucrose cushion centrifugation (50 mM Tris-HCl, 500 mM NaCl, 50 % sucrose, pH 7.0). The inclusion bodies were dissolved in denaturing buffer (6 M GdnHCl, 50 mM Tris-HCl, 100 mM NaCl, 20 mM imidazole, pH 7.5), applied to Ni-NTA agarose resin (Cytiva, Uppsala) and then VDAC2 was eluted with elution buffer (6 M GdnHCl, 50 mM Tris-HCl, 100 mM NaCl, 250 mM imidazole, 1mM TCEP, pH 7.5). Purified VDAC2 in elution buffer was precipitated by dialysis against 4 liters of dialysis buffer (50 mM Tris-HCl, 50 mM NaCl, 1 mM EDTA, 1 mM TCEP, pH 7.5) in a 12,000-14,000 molecular weight cutoff (MWCO) dialysis membrane. Precipitated VDAC2 was isolated by centrifugation at 12,000 g.

**Refolding and purification of VDAC2 mutants in LDAO detergent**

Purified precipitated VDAC2 was dissolved in guanidine hydrochloride buffer (100 mM NaPi, 100 mM NaCl, 6 M GdnHCl, 2 mM TCEP, 1 mM EDTA, pH 6.8) at a concentration of 2.5-3 mg/ml. VDAC2 was refolded at 4°C by dropwise dilution of one volume of VDAC2 solution into ten volumes of refolding buffer (100 mM NaPi, pH 7.0, 100 mM NaCl, 1 mM EDTA, 2 mM TCEP, 1.5% (64.5 mM) LDAO (Anatrace, Maumee) with stirring. The final ratio of VDAC2 protein to LDAO micelles is about 1:90 assuming the aggregation number of LDAO micelle is 76. After overnight stirring at 4℃ the refolded VDAC2 sample was dialyzed against 4 liters of buffer (25 mM NaPi, 1 mM EDTA, 1 mM TCEP, pH 7.0) for 12 to 16 hours. Little amount of precipitated VDAC2 was removed using centrifugation at 12,000 g, followed by filtration with a 0.2-μm membrane.

Cation exchange chromatography was employed to isolate properly folded VDAC2 protein in LDAO detergent micelles. The sample was loaded onto a 20 ml SP Sepharose HP column equilibrated with buffer A (25 mM NaPi, 1 mM EDTA, 1 mM TCEP, 0.1% LDAO, pH 7.0). VDAC2 was eluted during a 5%-55% gradient with buffer B (25 mM NaPi, 1 mM EDTA, 1 mM TCEP, 0.1% LDAO, 1M NaCl, pH 5.9) at about 35% buffer B. Pure VDAC-2 fractions were pooled and concentrated to the desired concentration using 30kDa molecular weight cutoff (MWCO) concentrators. VDAC2 was further purified by gel filtration chromatography using a Superdex 200 prep grade column (Cytiva, Wilmington DE), pre-equilibrated with NMR buffer (25 mM NaPi, 50 mM NaCl, 0.08% LDAO, 2 mM EDTA, pH 6.5).

**Channel reconstitution and conductance measurements**

Planar lipid membranes were formed on a ~ 70−100-μm diameter orifice in a 15-μm thick Teflon partition that separated two compartments filled with 1 M KCl buffered with 5 mM HEPES at pH 7.4, as described previously [1]. Lipid monolayers for membrane formation were made from 5 mg/mL solution of lipid in pentane. Single-channel data were filtered by a low-pass eight-pole Butterworth filter (Model 900 Frequency Active Filter; Frequency Devices, Ottawa, IL) at 15 kHz and saved with a sampling frequency of 50 kHz and analyzed using pClamp 10.7 Software (Molecular Devices, San Jose, CA). Sall data were collected at room temperature of 21.0 ± 1.0 ºC.

**Measuring of voltage-gating on multichannel membrane**

VDAC voltage-dependent gating was measured using a previously described protocol [2], in which gating is inferred from the response of multiple channels to a slow symmetrical 5 mHz triangular voltage wave of ± 60 mV amplitude applied with a 33220A arbitrary waveform generator (Agilent Technologies, Santa Clara, CA). VDAC multichannel recordings were filtered by a low-pass eight-pole Bessel filter at 1 kHz, digitized at a sampling frequency of 2 Hz, and analyzed, as described previously [2], using pClamp 10.7 Software and an algorithm developed in house [3].

**Supplemental Figures**

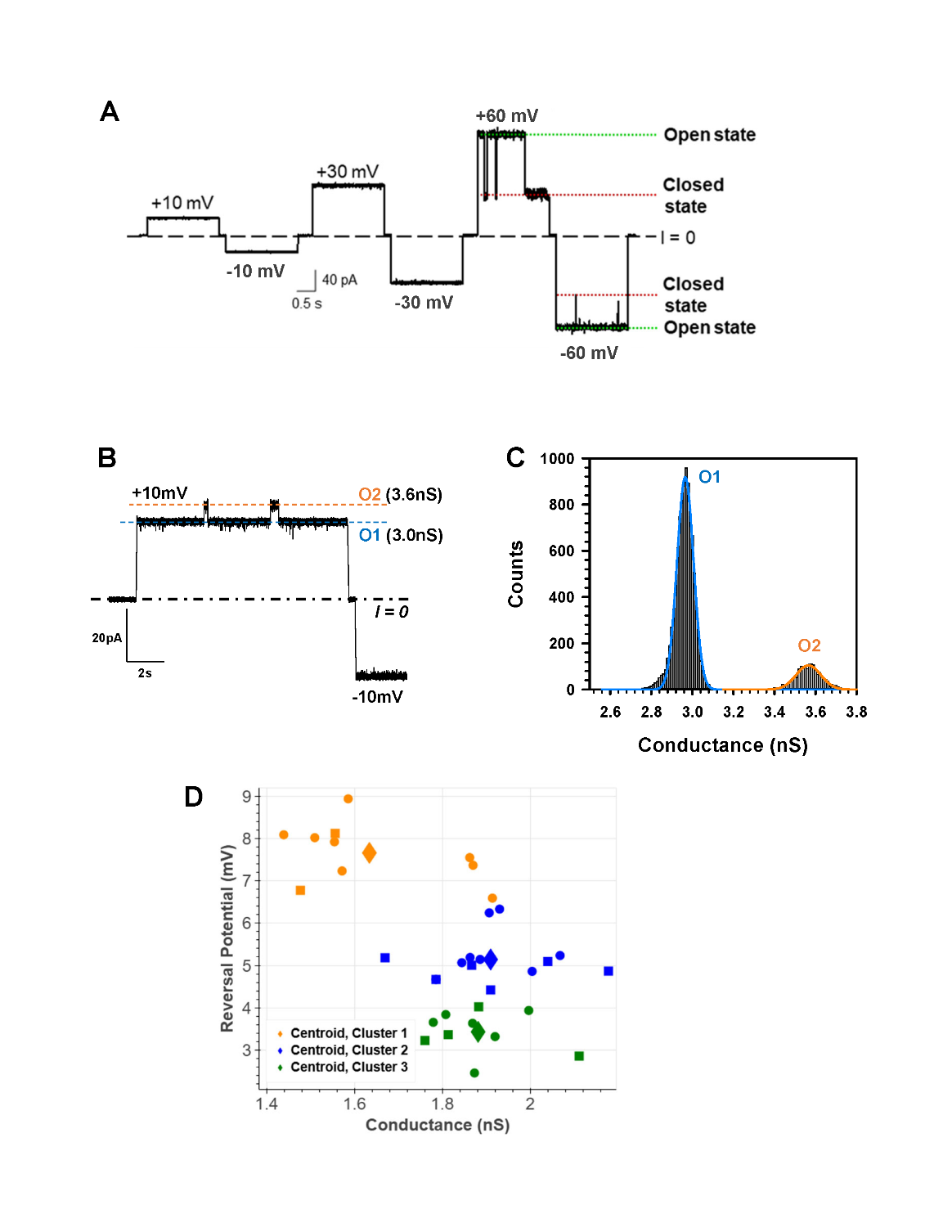

**Supplemental Figure S1. Conductance substates in VDAC2 WT are observed at low voltages.** (**A**) A representative current record of single VDAC2 at ±10 mV of applied voltage. Two substates O1 of 3.0 nS and O2 of 3.6 nS are indicated by blue and orange lines, respectively. The current record was digitally filtered at 500 Hz using a low-pass Bessel (8-pole) filter. (**B**) The conductance histogram corresponding to the current trace in (A) at + 10 mV shows two well-separated conductance levels. (**C**) Example of a VDAC2 displaying no substates across multiple applied voltages. (**D**) K-means clustering plot of Conductance vs Reversal Potential for VDAC2 WT obtained in a 5:1 KCl salt gradient. Each individual measurement is colored to the cluster in which it lies. Each of the three larger diamonds represents the mean of one of the calculated clusters. Planar lipid membrane was formed from 2PG/PC/PE lipid mixture in 1M KCl, pH 7.4.

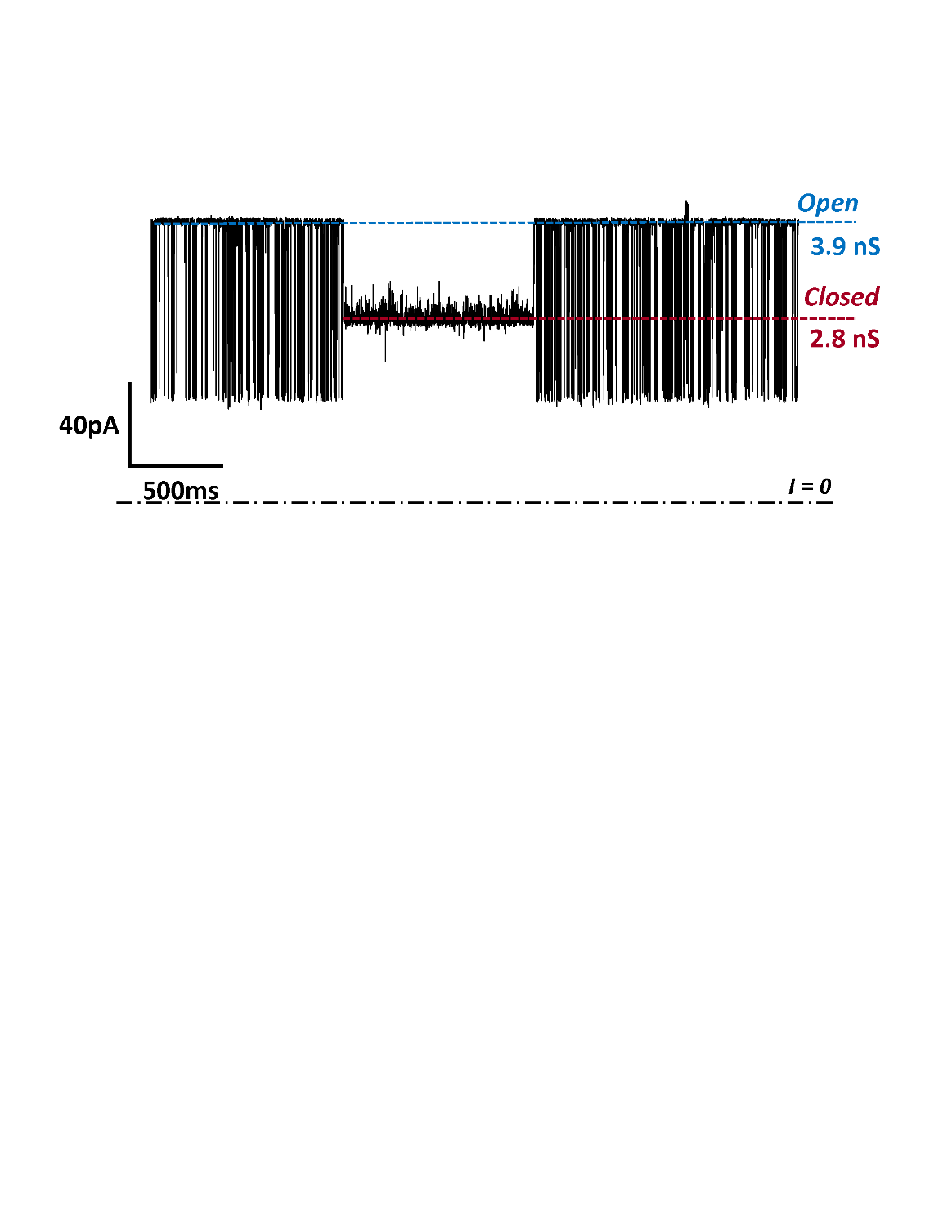

### **Supplemental Figure S2.** **The voltage-induced low conducting, closed state of VDAC2 is not blocked by αSyn**. The applied voltage was +32.5 mV. 10 nM αSyn on both sides of the membrane. The current record was digitally filtered using a 1 kHz low-pass Bessel (8-poles) filter. The planar membrane was formed from 2PG/PC/PE in 1 M KCl at pH 7.

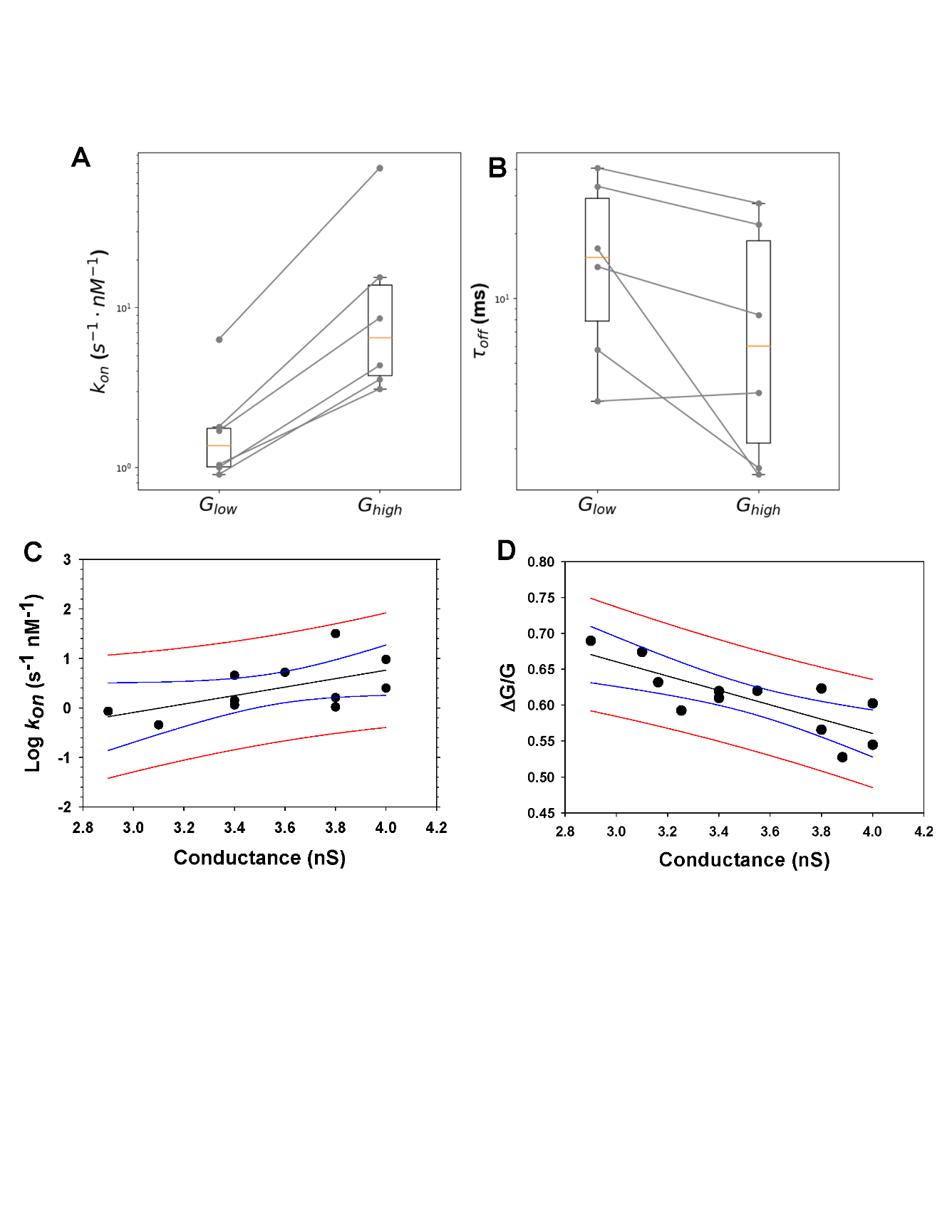

**Supplemental Figure S3. Statistical analysis of the kinetic parameters of αSyn interaction with different VDAC2 conductance states.** Box and Whisker plot for *kon* (**A**) and *τoff* (**B**) of higher conducting *GHigh* and lower conducting substates *GLow* observed in independent single-channel experiments with VDAC2 WT. (**C, D**) Lines represent the statistical analysis of the linear correlation between the on-rate (*kon*) of αSyn-VDAC2 WT interaction (C) and the conductance of αSyn-blocked state (*ΔG/G*) (D) with the conductance of different substates (N=12) in 7 individual channels recorded in different PLM. *ΔG/G* is a relative conductance, where *ΔG* is the difference between open (*G*) and blocked state conductances for each substate. Black lines are linear regressions with R2 = 0.33 (A) and 0.62 (B). Blue lines represent 95% of the confidence band, and red lines represent 95% of the prediction band. Data are obtained at 32.5 mV. All experimental conditions are as in Fig. S2. Statistical analysis of the substates was performed using the nonparametric paired Wilcoxon signed rank test, comparing substates observed on the same channel, allowing comparison between data taken at different voltages.

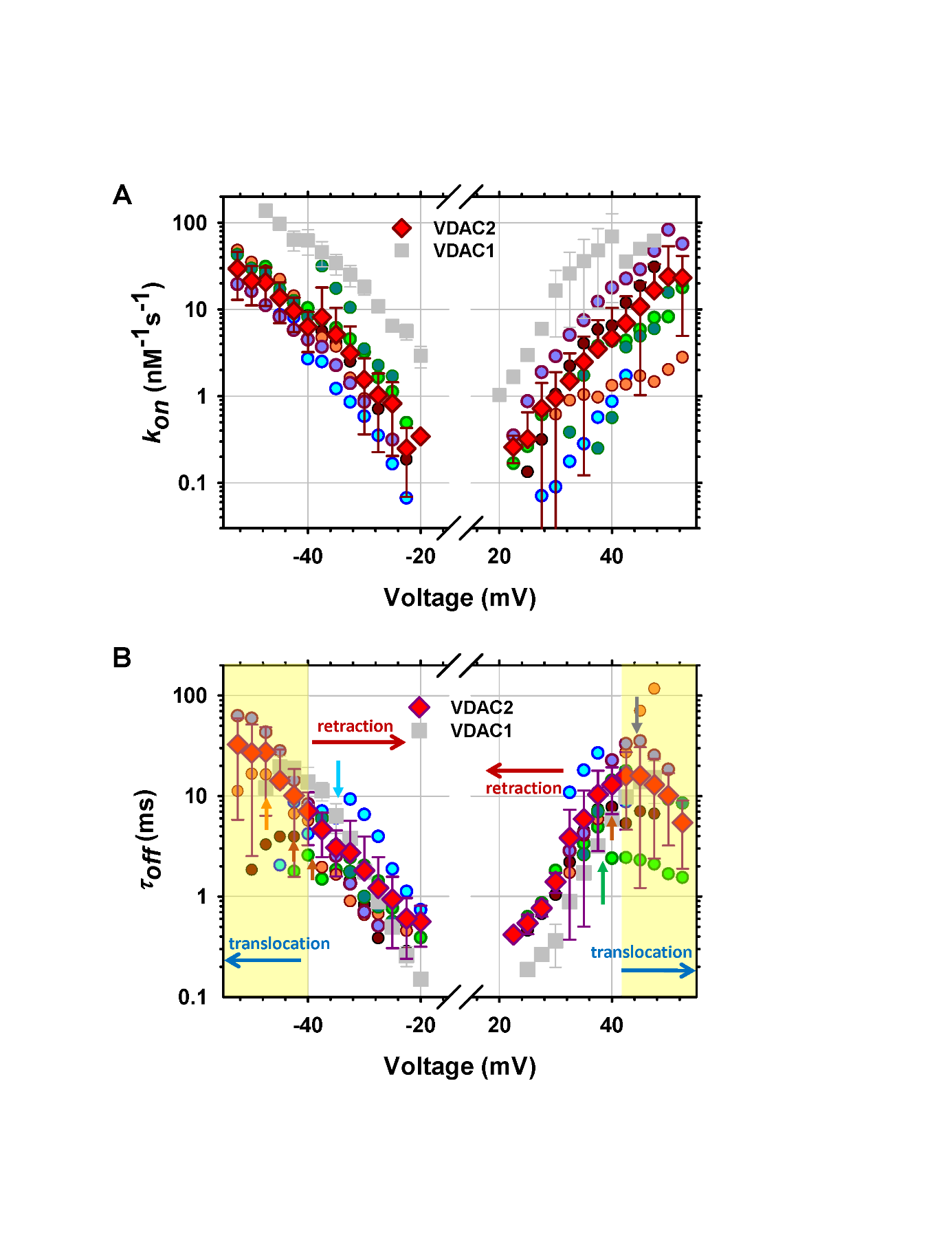

**Supplemental Figure S4. Voltage dependences of the kinetic parameters of αSyn-VDAC2 interactions.** Voltage dependences of *kon* (**A**) and blockage time, *τoff*, (**B**) obtained in 7 individual VDAC2 WT channels in the presence of 10 nM αSyn in both compartments (circular symbols). Each color of the circular symbol represents a separate experiment. The analyzed conductances vary from 2.9 to 4.1 nS with an average of 3.5± 0.4 nS. Red diamond symbols are means ± SD of 7 experiments. Grey squares are mean data for VDAC1 WT ± SD (N=4) shown for comparison. (A) - Despite high variability in *kon* between channels and substates, all *kon* for VDAC2 WT are below *kon* values for VDAC1. (B) - The increase of *τoff* with voltage amplitude corresponds to the αSyn blockage/retraction regime, which is similar for individual VDAC2 channels and for VDAC1. The decrease of *τoff* corresponds to the translocation regime indicated by yellow highlights. The voltages at which the blockage/retraction regime becomes translocation vary from -32.5 to -52.5 mV at negative polarities and from 37.5 to 45 mV at positive and indicated by arrows. In some experiments, the translocation regime starts at *V* > |55 |mV. Other experimental conditions, as in Fig. S2.

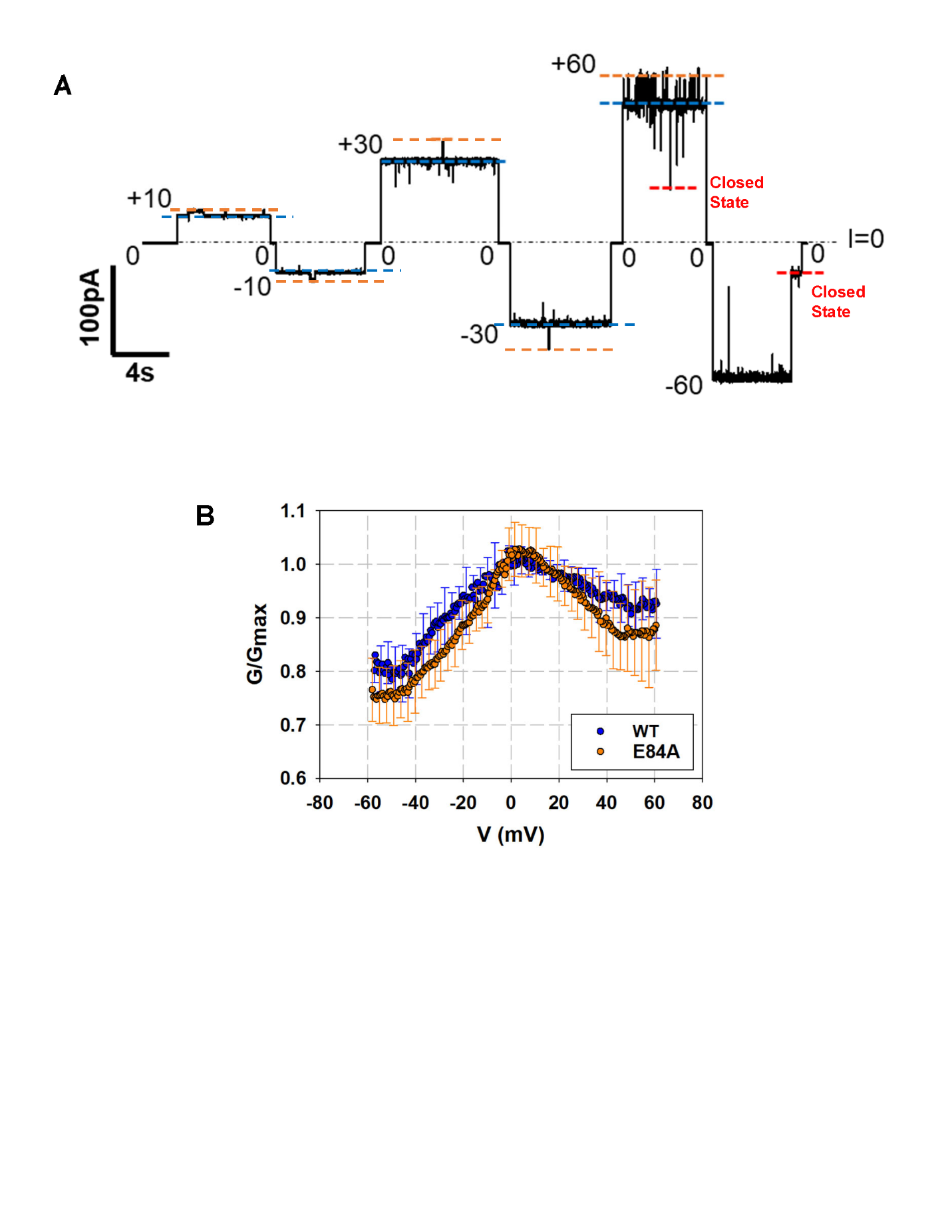

**Supplemental Figure S5.** **VDAC2 E84A mutant forms a functional voltage-gating channel with multistate behavior.** (**A**) -Representative single-channel current traces obtained with reconstituted VDAC2 E84A inserted into 2PG/PC/PE membrane in 1M KCl at pH 7.4 demonstrates one main conducting state of 3±0.1nS (blue dashed lines) and higher conducting substate of 3.5 ± 0.1 nS (orange dashed lines) at all applied voltages starting from ±10 mV. The dash-dotted line represents zero current level; dashed lines indicate conductance substates. Channel is characteristically voltage-gated – transition to the lower conducting state of (red dashed line) at - 60mV. The current record was digitally filtered at 500 Hz using a digital low-pass Bessel (8-pole) filter. (**B**) - Characteristic bell-shaped plots of normalized average conductance as functions of the applied voltage for WT and E84A VDAC2. *G*/*G*max is the normalized conductance, where *G*max is the maximum conductance at voltages closest to 0 mV. Data are means of 3-4 independent experiments ± S.D.

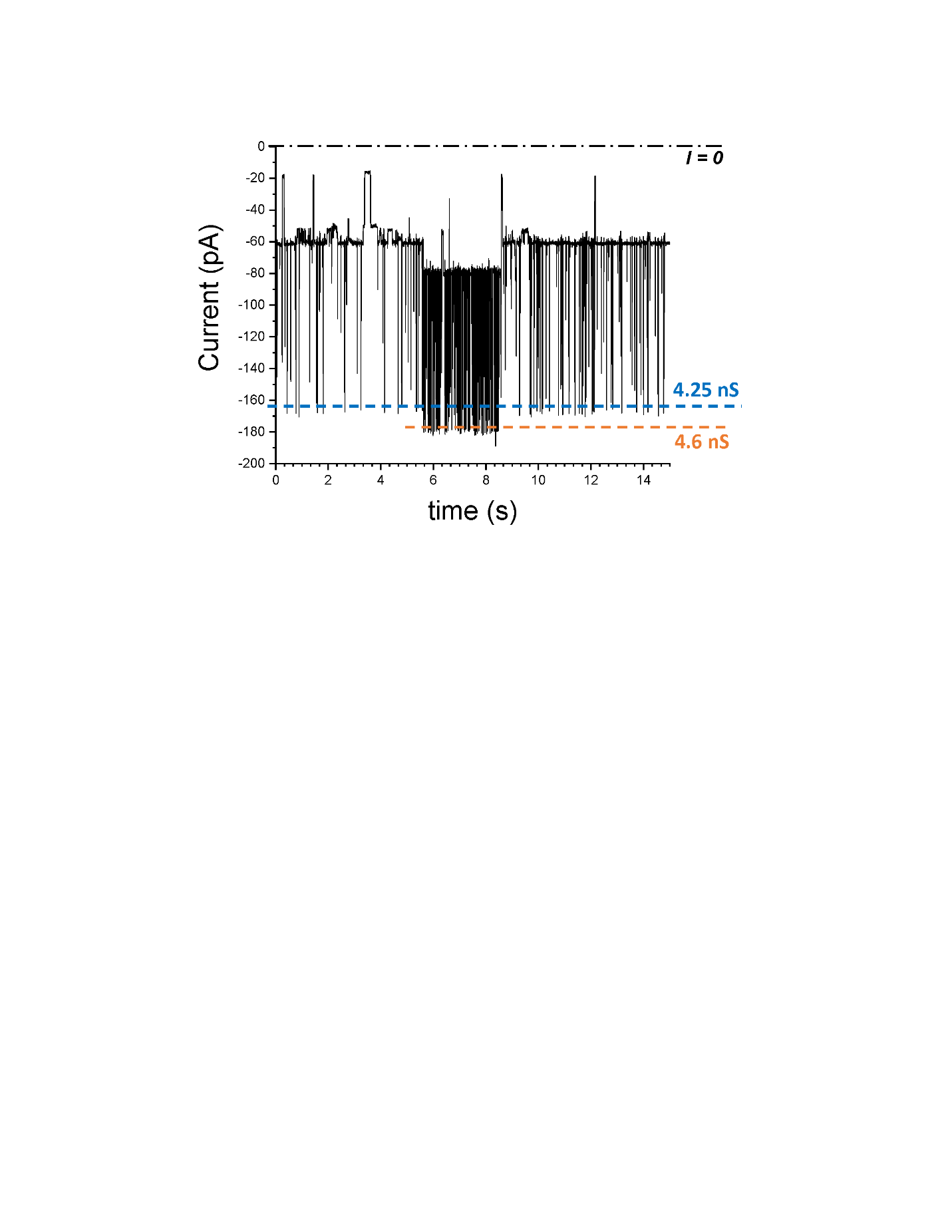

**Supplemental Figure S6.** **Zebrafish VDAC2 (zfVDAC2) has conductance substates that interact differently with αSyn**. The representative current record of single zfVDAC2 channel reconstitute to the PLM formed from DPhPC in 1M KCL, 5mM HEPES, pH 7.4 in the presence of 10 nM of αSyn added to both sides of the membrane at – 40 mV of applied voltage. Channel conductance transitions between two levels of 4.25 and 4.6 nS. The intensity of blockage events is higher for the higher conductance level. The trace was digitally filtered at 1 kHz.
